# Functional organization of the spinal locomotor network based on analysis of interneuronal activity

**DOI:** 10.1101/2025.05.06.652406

**Authors:** Pavel E. Musienko, Oleg V. Gorskii, Tatiana G. Deliagina, Pavel V. Zelenin

## Abstract

Locomotion is a vital motor function for any leaving being. In vertebrates, a basic locomotor pattern is generated by the spinal locomotor network (SLN). Although SLN has been extensively studied, due to technical difficulties, most data were obtained during fictive locomotion, and data about activity of spinal neurons during locomotion with intact sensory feedback from limbs are extremely limited. Here, we overcame the technical problems and recorded activity of putative spinal interneurons from spinal segments L4-L6 during treadmill locomotion (with intact sensory feedback from limbs) evoked by stimulation of the mesencephalic locomotor region in the decerebrate cat. We analyzed activity phases of recorded interneurons, by using a new method that took into account the previously ignored information about stability of neuronal modulation in the sequential locomotor cycles. We suggested that neurons with stable modulation (i.e. small dispersion of their activity phase in sequential cycles) represent the core of SLN. Our analysis allowed to reveal functional groups of neurons with stable modulation presumably generating the vertical and horizontal components of the step, and to characterize their location in the spinal cord. Analysis of relationships between activity phases of these groups revealed possible connections between them, suggesting a novel model for generation of locomotion that combines reciprocally active half-centers with a ring consisting of four sequentially active groups, each inactivating the preceding one and activating the next one.

## 1 INTRODUCTION

For most legged vertebrates, forward locomotion represents the main form of progression, although they can also perform backward, sideways and in place stepping. In all studied vertebrates, including humans, the basic pattern of stepping movements in any direction is generated by a neuronal network residing in the spinal cord (the spinal locomotor network, SLN). SLN includes a network capable of generating the basic locomotor pattern without the sensory feedback (Brown, 1911) currently termed as “the central pattern generator”, CPG. However, normally this basic pattern is substantially modified by the activity of the sensory neurons that constitute an important part of SLN (Rossignol et al., 2006). The descending commands from the brain turn SLN on and off, control the speed and gait of locomotion, and modify the basic pattern for steering, obstacle avoidance, etc. (Orlovsky et al., 1999; Bouvier et al., 2015; Caggiano et al., 2018). Forward locomotion can be evoked, and its speed and gait can be controlled in the decerebrate preparation by stimulation of the mesencephalic locomotor region (MLR) of the brainstem (Shik et al., 1966; Shik & Orlovsky, 1976). This experimental paradigm has been extensively used for investigation of SLN. The results of these experiments together with studies of the locomotor kinematics and the motor pattern in intact and spinal animals (Halbertsma, 1983; Krouchev et al., 2006; Markin et al., 2012) have led to the development of computational models of SLN that include interneurons, motoneurons, limb dynamics and sensory feedback (McCrea & Rybak, 2008; Rybak et al., 2015; Shevtsova et al., 2015; Markin et al., 2016). However, due to technical difficulties related to the recording of spinal neurons during locomotor movements, the experimental data related to operation of SLN during “real locomotion” that is locomotion with close to normal sensory feedback, are extremely limited (Orlovsky & Feldman, 1972; Feldman & Orlovsky, 1975). Also, spinal interneurons were recorded during forward air stepping, when the movement-related sensory feedback was present but substantially different from the normal one (AuYong et al., 2011).

Most data related to activity of spinal interneurons were obtained during fictive locomotion in immobilized animals or in *in vitro* newborn mice preparations. Thus, activity of identified spinal interneurons (Renshaw cells and Ia interneurons) was recorded during fictive locomotion caused by stimulation of MLR (Noga et al., 1987; Pratt & Jordan, 1987; Shefchyk et al., 1990). Since it was demonstrated that MLR is a center for initiation of forward locomotion only (Musienko et al., 2012), most likely, these neurons were recorded during fictive forward locomotion. Also, activity of non-identified spinal interneurons was recorded during spontaneous episodes of fictive locomotion in paralyzed cats (Baev et al., 1979) and in anesthetized rabbits (Viala et al., 1991). In these studies, as well as in studies on *in vitro* preparations (Zhong et al., 2012), the neuronal activity was recorded along with activity of motor nerves only to a pair of antagonistic muscles in each limb, and thus, it was impossible to determine if forward, backward or in place fictive locomotion was generated. Our earlier results suggest that SLNs generating forward and backward stepping differ to some extent (Merkulyeva et al., 2018; Musienko et al., 2022). Thus, it is difficult to assess whether the models based on data obtained during fictive locomotion represent operation of SLN during real forward locomotion.

The aim of the present study was to analyze operation of SLN during real forward locomotion with normal sensory feedback from limbs. For this purpose, we recorded activity of putative spinal interneurons during real locomotion evoked in decerebrate cat by MLR stimulation. We employed a new approach for analysis of locomotion-related activity of neurons. Our analysis allowed to reveal functional groups of neurons, to suggest functional connections between them, to characterize their distribution in the spinal cord and to propose a novel model of SLN for generation of forward locomotion.

A brief account of this study was published in abstract form (Zelenin et al., 2019).

## 2 MATERIALS AND METHODS

### 2.1 Animals

Experiments were carried out on five adult cats of either sex (weighing 2.5-4.0 kg). The same animals were used in our previous study (Musienko et al., 2020). All procedures were conducted in accordance with a protocol approved by the Animal Care Committee of the Pavlov Institute of Physiology, St. Petersburg, Russia (protocol #01a/2016), and followed the European Community Council Directive (2010/63EU) and the guidelines of the National Institute of Health *Guide for the Care and Use of Laboratory Animals*.

### 2.2 Surgical procedures

The surgical procedures were the same as in our previous study (Musienko et al., 2020). The cats were deeply anesthetized with isoflurane (2-4%) delivered in O_2_. The trachea was cannulated, and carotid arteries were ligated. The animals were decerebrated at the precollicular-postmammillar level. A laminectomy was performed in the lumbar area. Anesthesia was discontinued after the surgical procedures, and the experiments were started 2–3 h thereafter. During the experiment, the rectal temperature and mean blood pressure of the animal were continuously monitored and were kept at 37±0.5°C and >80 mmHg.

### 2.3 Experimental design

The experimental design (Figure 1A) was similar to that used in the previous study (Musienko et al., 2020). The head, the vertebral column, and the pelvis of the decerebrate cat were fixed in a rigid frame (Figure 1A). The forelimbs had no support, whereas the hindlimbs were positioned on a treadmill with two separate belts (left and right moving backward in relation to the animal at the same speed (0.5 m/s) and below referred to as the “treadmill belt”. The distance between the treadmill belt and the fixed pelvis was 21–25 cm (depending on the animal’s size), which determined a semi-flexed limb configuration in the middle of stance typical for walking.

**FIGURE 1.**
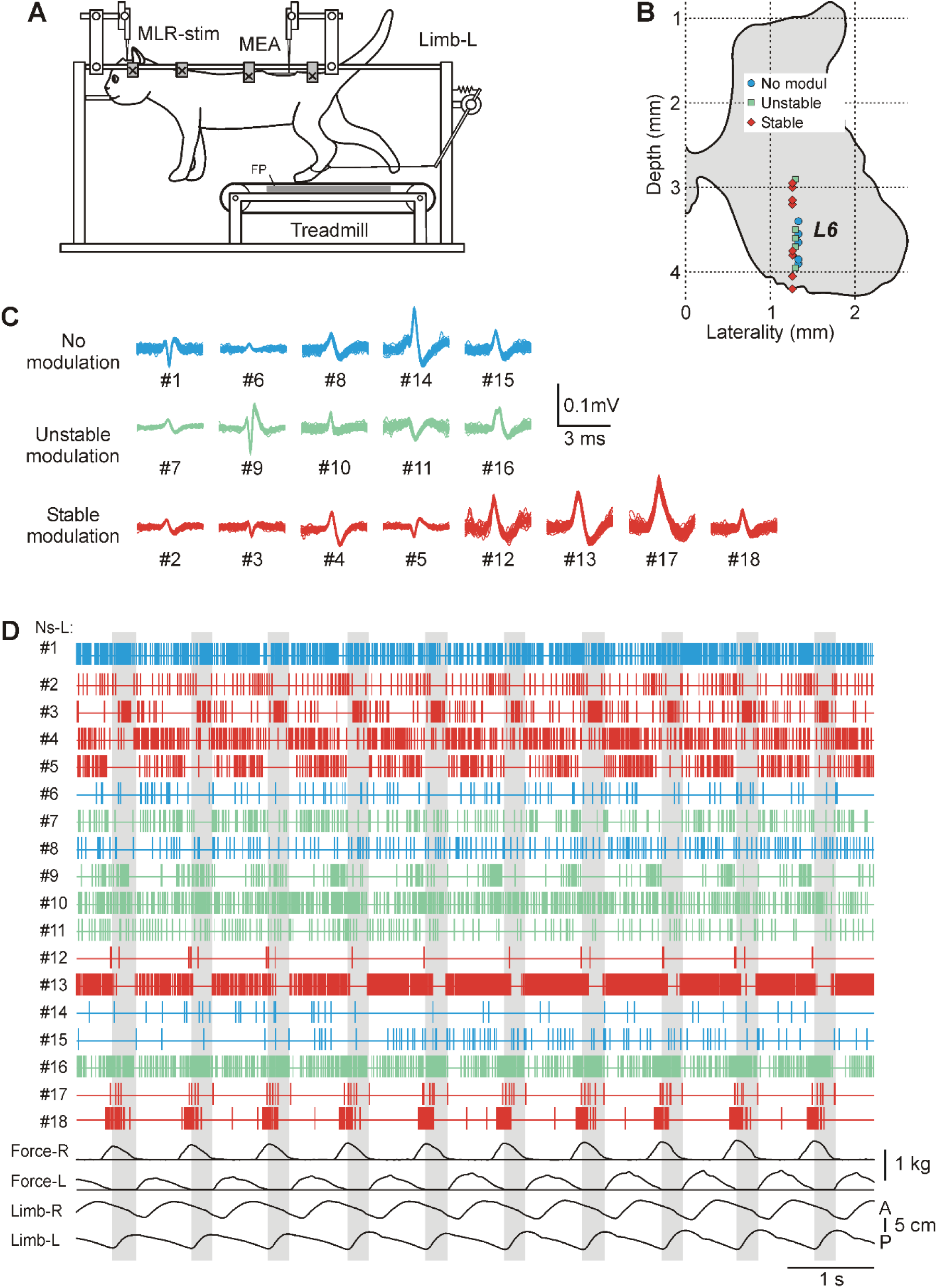
Recording of the activity of spinal neurons during MLR-evoked locomotion. (A) Experimental design. MLR-stim, an electrode for stimulation of mesencephalic locomotor region. MEA, a microelectrode array for recording neuronal activity. Limb-L, a mechanical sensor recording anterior/posterior movements of the left hindlimb (a sensor recording the right hindlimb movements is not shown). FP, a force plate. (B-D) An example of 18 spinal neurons recorded simultaneously in L6 during locomotion. Red and green, neurons with stable and unstable modulation, respectively. Blue, non-modulated neurons. (B) Locations of the neurons. (C) Superimposed spikes of individual neurons extracted from the mass activity. Recording depths (D) are indicated. (D) Activity of individual neurons (##1-18) during locomotion recorded along with hindlimb movements (Limb-R, Limb-L) and contact forces (Force-R, Force-L). Swing phases of the left hindlimb are highlighted.

Locomotion was evoked by electrical stimulation of the mesencephalic locomotor region (MLR) (Shik et al., 1966; Shik & Orlovsky, 1976; Jordan, 1986; Garcia-Rill & Skinner, 1987a,b). The stimulation started in 2–3 s after the onset of the treadmill belt motion. For MLR stimulation, a bipolar electrode (two 150 μm wires isolated except for the tips and separated by 0.5 mm) was inserted into the brainstem area (Horsley-Clarke coordinates P2, R/L4, H0) by means of a micromanipulator (MLR-stim in Figure 1A). We used the following parameters of stimulation: frequency, 30 Hz; pulse duration, 0.5-1 ms; current, 50-200 µA.

We recorded the anterior-posterior locomotor limb movements by means of two mechanical sensors (one of which, Limb-L, is shown in Figure 1A), as well as the vertical forces developed by each of the limbs by means of two force plates positioned under the left and right parts of the moving belt (FP in Figure 1A).

### 2.4 Electrophysiology

Neurons were recorded extracellularly by means of commercially available multichannel electrode arrays (MEA in Figure 1A). Each array consisted of a shaft with 32 Pt/Ir/Au electrode sites (A1x32-10mm-50-177; Neuronexus, Ann Arbor, MI). Each site surface area was 177 µm^2^. Electrode sites were distributed 50 µm apart vertically, and thus the recording length of the array was approximately 1.5 mm, which allowed for simultaneous recordings of many individual neurons from different areas of the gray matter (Figure 1B). The array was mounted on a custom electrode holder driven by a manual micromanipulator. We attempted to systematically explore the whole cross-section of the gray matter except for the motor nuclei areas. Activity of putative spinal interneurons during MLR-evoked locomotion was recorded along with ground reaction forces and signals from mechanical sensors in ∼20-60 locomotor cycles. Only episodes with stable well-coordinated locomotion (similar limb trajectories, stable forces, the right and left hindlimbs stepping in antiphase) were analyzed. For all analyzed episodes, the relative variability (SD/mean) of the cycle duration was <12%, the relative variability of the swing proportion <17%, the relative variability of the step length <20%, the right and left hindlimb movements were close to anti-phase (deviation of the inter-limb phase shift from 0.5 was <0.05).

To increase the stability of recording that is to prevent displacement of the nervous tissue in relation to the recording electrodes caused by movements (locomotor, breathing, etc.), the space between the dura matter and walls of the spinal canal was filled by tissue adhesive glue Indermil X Fine (Henkel Ireland Operations and Research Ltd, Dublin, Ireland) in the area of recording.

Signals from the electrode array (neuronal activity), from mechanical sensors and force plates were amplified, digitized with a sampling frequency of 30 kHz (neurons) and 1 kHz (sensors), and recorded on a computer disk using a data acquisition and analysis system (Power1401/Spike2, Cambridge Electronic Design, Cambridge, UK). The multiunit spike trains recorded by each electrode were separated into unitary waveforms representing the activity of individual neurons using the spike-sorting procedure in the same system. The spikes with the same waveform were supposedly generated by the same spinal neuron during locomotion (Figure 1C). Only neurons with a stable spike shape were used for analysis. Many neurons were recorded simultaneously by several neighboring sites, which increased the confidence of the spike sorting. An example of the activity of 18 neurons simultaneously recorded during locomotion, the waveforms of their spikes, and positions of the neurons on the cross-section of the gray matter are shown, respectively, in Figures 1, D, C, and B. To exclude recording of the same neuron twice, medio-lateral distance between recording tracks was at least 0.2 mm (while the distance between recording sites that could pick up spikes of the same neuron was not more than 0.1 mm).

### 2.5 Analysis of activity of individual neurons

Neurons with the mean firing frequency less than 1 Hz were considered inactive during locomotion (Table 1). The activity of neurons was typically modulated in the rhythm of stepping movements (Figure 1D). To characterize this modulation, the phase histogram of neuronal activity in a step cycle of the ipsilateral limb was generated. Because of some variability in the duration and structure of step cycles within a test and between tests in different animals, we used dual-referent phase analysis (Berkowitz & Stein, 1994), that is we normalized swing and stance phase separately. Step cycle durations were normalized to unity. Swing and stance phases were normalized, respectively, to 0.38 and 0.62 parts of the locomotor cycle that represent proportions of swing and stance in the locomotor cycle averaged across all episodes of locomotion in all cats (for details, see Supplementary Data, page 3). Such normalization ensured that neuronal activity during a definite phase (swing or stance) during one walking episode was compared to activity during the same phase in the other episodes, or when these characteristics were compared in different steps within the same episode.

**TABLE 1.** Descriptive statistics for the recorded population. *Spinal segment*, the segment in which neurons were recorded in a particular cat. The number of neurons as well as the percentage of neurons out of entire population of neurons recorded in an individual cat, or in a segment, or of the entire population of recorded neurons are indicated in parenthesis.

| Cat # | Spinal segment | Recorded neurons | Not active | Non-modulated | Modulated neurons | Stable neurons |
| --- | --- | --- | --- | --- | --- | --- |
| 1 | L4 | 91 (100%) | 2 (2%) | 11(12%) | 78 (86%) | 48 (53%) |
| 2 | L4 | 9 (100%) | 0 | 1(11%) | 8 (89%) | 7 (77%) |
| 3 | L4 | 28 (100%) | 1 (3%) | 3(11%) | 24 (86%) | 15 (54%) |
| Total | L4 | 128 (100%) | 3 (2%) | 15(12%) | 110 (86%) | 70 (55%) |
| 4 | L6 | 35 (100%) | 1 (3%) | 2 (6%) | 32 (91%) | 24 (69%) |
| 5 | L6 | 80 (100%) | 6 (7%) | 23 (29%) | 51 (64%) | 32 (40%) |
| Total | L6 | 115 (100%) | 7 (6%) | 25 (22%) | 83 (72%) | 56 (49%) |
| Total | L4+L6 | 243 (100%) | 10 (4%) | 40 (17%) | 193 (79%) | 126 (52%) |

Typically, neurons discharged only during a part of the step cycle producing one burst of activity. Some neurons had their bursts on top of the background tonic activity. To estimate the onset and offset phases of the neuron’s burst in the step cycle, the spike time sequence during one step (Figure 2A) was converted to the instantaneous rate versus time (Figure 2B) and then to instantaneous rate versus phase (1000 data points per normalized cycle; Figure 2C). The resulting curve was approximated with the best two-level rectangular fit (shown by red lines in Figure 2C; Zelenin et al., 2011). This fit provided the burst onset phase and offset phase expressed as a proportion of the normalized cycle (Figure 2C; for details, see Supplementary Data, page 4). The phases (Figure 2D) were averaged across all step cycles using the circular statistics methods (Batschelet, 1981), providing the mean onset and offset phases, as well as the corresponding standard deviations, characterizing stability/variability of the phases (Figure 2E; for details, see Supplementary Data, page 5).

**FIGURE 2.**
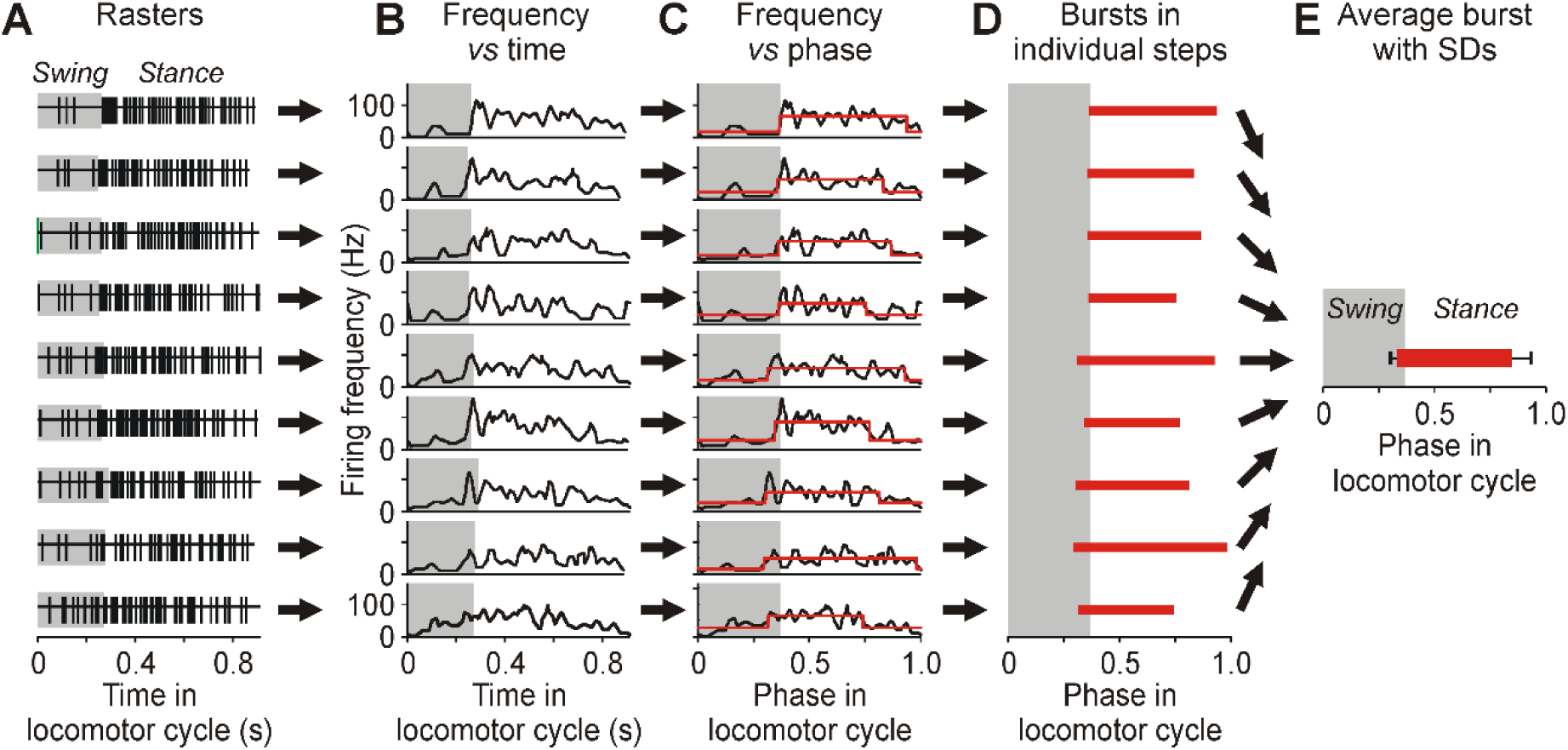
Analysis of the activity of spinal neurons during MLR-evoked locomotion. (A-E) An example of analysis of activity (neuron #4 from Figures 1B-D). For each individual step, a spike raster aligned at the swing start (A), the instantaneous firing rate *vs* time (B), the rate *vs* phase with dual-reference phase normalization (C), and the burst phase (D) were determined. Burst phases in individual steps were used to calculate the average and *SD* for burst on- and offset phases (E).

We supposed that neurons with stable phase of activity have a larger contribution to control of locomotor movements. Stability of the phase is characterized by its *SD*: the smaller *SD* reflects the higher stability. We proposed one novelty in analysis: assigning more weight to the data with lower variability (smaller *SD*). With this purpose, we converted *SD*s into “weights” by using a formula: *w* = 10 / (1+900*SD*^2^) (for details, see Supplementary Data, page 6). Thus, neurons with higher variability (that is neurons with unstable activity, and therefore larger *SD*) had lower weight of their phase, and *vice versa*. To take into account both the onset stability and the offset stability of a neuron, we calculated the integral stability of the activity phase of the neuron during locomotion (*W*): *W* = (*w*_on_)^2^ + (*w*_off_)^2^, where *w*_on_, *w*_off_ were the weights of the onset and offset phases, respectively.

Averaging of the instantaneous rate versus phase across all recorded steps produced the mean phase profile of the neuron’s activity. A neuron was considered modulated if the activity profiles in individual steps were similar to the average profile (the Pearson’s correlation coefficient was higher than 0.3 in more than 60% of the locomotor cycles). The other neurons, with more random bursts of activity, were considered non-modulated (Table 1; blue neurons in Figures 1B-D).

Modulated neurons were further classified into two subpopulations: with stable modulation and with unstable modulation. We used two criteria for stability of modulation of the neuron. (i) SD of the burst onset and/or burst offset had to be not more than 0.1 part of the locomotor cycle. We accepted stability of only one edge of the burst since some of the recorded neurons had a ramp-up or ramp-down burst shape with stable burst offset and burst onset phase, respectively. (ii) The average coefficient of correlation between the profiles of the activity in individual locomotor cycles and in the entire activity histogram should not be less than 0.6. Modulation was considered stable if both criteria were satisfied. Examples of neurons with stable and unstable modulation (their locations, spike shapes, and activity) are shown in Figures 1B-D (red and green, respectively).

It should be noted that we analyzed activity of neurons in relation to the locomotor cycle of the ipsilateral limb. However, we cannot exclude that some of the recorded neurons were commissural interneurons contributing to control of the contralateral limb.

### 2.6 Analysis of population distribution of burst onset and offset phases

The population distribution of any measurable parameter (e.g., the burst offset phase measured in a set of neurons) is traditionally presented with a distribution histogram. We proposed a modified presentation that took into account not only the measured values but also the precision of these measurements. Specifically, each phase *φ_i_* measured with a precision *SD_i_*, was presented with the probability density *p_i_(φ)*: a Gaussian distribution with the mean value equal to *φ_i_* and the standard deviation equal to *SD_i_*. The average of such distributions across the entire population of neurons provided the probability density to observe a particular value of the measured parameter. Besides taking into account the precision of measurements, this procedure also smoothens the resulting curves. Analogous to the described above probability distribution of onset/offset phases, the weighted probability distribution of onset/offset phases was calculated. With this purpose the weighted averaging was applied to the distributions for individual neurons: Σ(*w_i_p_i_*(φ)) / Σ*w_i_* where *w_i_* was the weight function described in Matearials and Methods 2.5. For details of these procedures, see Supplementary Data, page 7.

### 2.7 Cluster analysis

To reveal groups of neurons with similar activity phases, we used a modified version of the “nearest neighbor” method of cluster analysis similar to the method used by Krouchev et al. (2006). For each neuron, its activity phases were presented as a scatterplot in which the burst offset phase of a given neuron was plotted *vs* its burst onset phase (Krouchev et al., 2006) together with the onset and offset weights (Supplementary Figure 2A).

We did not specify the number of clusters *a priori*. The clustering procedure consisted of three stages: (i) ranking neurons according to stability of their activity, (ii) primary clustering (“seeding”) that produced an initial set of “raw” clusters, (iii) “refining” of the ”raw” clusters.

First, we ranked all neurons according to the integral stability of the activity phase of the neuron during locomotion (*W*, described in Matearials and Methods 2.5). Then, for primary clustering, the point in the scatterplot corresponding to the first neuron in the ranked list (characterizing the onset and offset of the neuron’s burst) was considered as the first cluster. Then points of other neurons were one by one compared with already existing clusters. Closeness or remoteness of a point from the centers of the existing clusters was characterized by two parameters: Euclidian distance (*D*) and Z-distance (*Z*). If at least one of these distances exceeded the threshold level (*D*>0.15 or *Z*>3) the point gave rise to a new cluster. If the point was close to one or several clusters (*D*<0.15 and *Z*<3), it was merged with the cluster for which *D* was minimal. For this expanded cluster, its center was re-calculated. The center of the cluster - that is the average onset and offset phases for neurons composing the cluster - was calculated by using the weighted average for a phase of the onset (Φ*^on^*) or offset (Φ*^off^*): Φ = Σ(*w_i_φ_i_*) / Σ*w_i_*, while the corresponding standard deviations were calculated with a formula: SD = Σ(*w_i_SD_i_*) / Σ*w_i_*.

The set of clusters obtained during the primary (“seeding”) clustering was used to re-sort the points with a “refining” procedure, similar to “seeding” but by using stricter similarity/closeness criteria: the point was considered far from the center of the cluster if *D*>0.1 or *Z*>3. In such a case it was considered as “assorted” (added to cluster 0). If the point was close to one or several clusters (*D*<0.1 and *Z*<3), it was merged with the cluster for which *D* was minimal. This resorting was repeatedly applied to the entire ranked list of neurons until convergence to the final set of *Clusters*. In addition, for each cluster the midburst phase Φ*^mid^* was calculated as Φ*^mid^* = (Φ*^on^* + Φ*^off^*) / 2.

Detailed description of the clustering analysis is presented in Supplementary Data (pages 8,9) and Supplementary Figures 2,3.

### 2.8 Statistical analyses

All quantitative data in this study are presented as mean±*SD*. To evaluate the statistical significance of difference in percentages of different *Clusters* of modulated neurons recorded in L4 and in L6, we used Fisher’s exact test; the significance level was set at *P* = 0.05.

### 2.9 Histological procedures

At termination of the experiments, cats were deeply anesthetized with isoflurane (5%) and then perfused with isotonic saline, followed by a 10% formalin solution. Frozen spinal cord sections of 50 µm thickness were cut in the regions of recording. The tissue was stained for Nissl substance with cresyl violet. The positions of the array in the spinal cord were verified by observation of the array track. Positions of recording sites were estimated in relation to the array track position.

## 3 RESULTS

### 3.1 Neuronal database

Altogether, the activity of 243 individual spinal neurons was extracted from the multiunit spike trains recorded in five decerebrate cats during treadmill forward walking evoked by MLR stimulation, including 128 neurons recorded in L4 and 115 neurons – in L6 (Table 1). Of these, few (4%) neurons were “inactive” (i.e. they had the mean cycle frequency of less than 1 Hz) and 17% were active but non-modulated (see criteria for modulation in Materials and Methods).

The majority of active neurons (∼86% from L4 and ∼72% from L6) were modulated in the rhythm of stepping movements [Table 1, Figure 1D (red and green neurons)]. Only these 193 modulated neurons were used for further analysis. Phase of bursts of individual modulated neurons recorded in L4 and L6 segments in the normalized locomotor cycle is shown in Figure 3. The neurons were ranked according to the value of the parameter “integral stability of the activity phase” during locomotion (*W*; see Materials and Methods). One can see that neurons with high values of *W* (with small *SD* of the burst onset and/or burst offset) located in the upper part of L4 and L6 neurons as well as neurons with low values of *W* (with large *SD* of the burst onset and burst offset) located in the lower part of L4 and L6 neurons had various phases of activity in the locomotor cycle.

**FIGURE 3.**
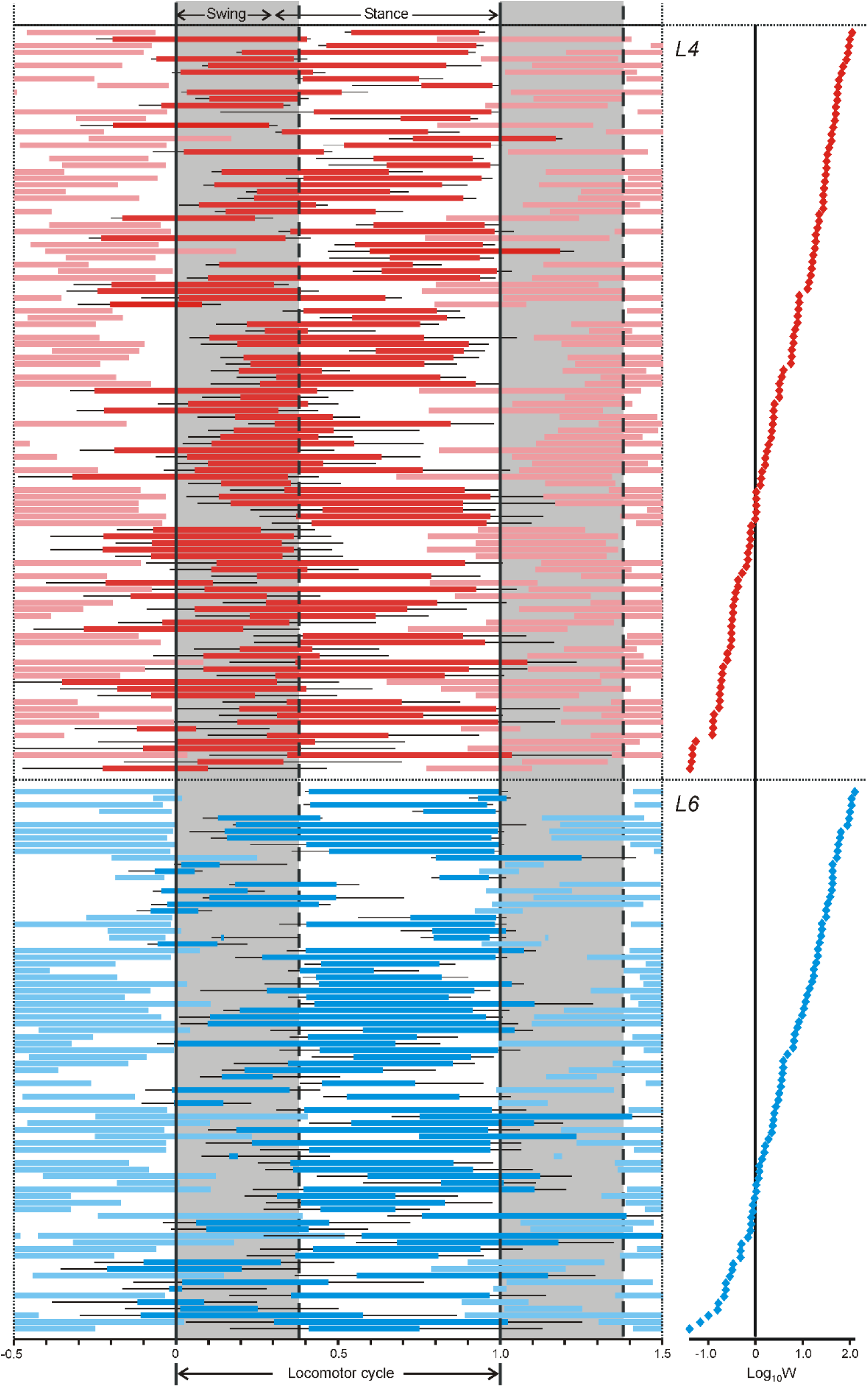
Phase distribution of bursts of individual neurons recorded in L4 and L6 segments in the normalized locomotor cycle. Neurons recorded in each segment are ranked according to the value of their parameter *W* (integral stability of the activity phase) shown on the right panel in logarithmic scale. Bursts of individual neurons recorded in L4 and L6 are indicated by thick red and blue lines, respectively. Activity of the same neurons in the preceding and the following cycles is indicated by the corresponding pale colors. Thin black lines indicate standard deviations of the burst onset and offset.

### 3.2 Distribution of burst onset and offset phases across the locomotor cycle

One can expect that phases of the locomotor cycle at which the burst onsets and burst offsets occur more frequently, represent the most critical/important phases for generation of the locomotor pattern (Orlovsky and Feldman, 1972). To reveal these key phases, the measured mean burst onset and burst offset phases of individual modulated neurons were used to calculate the probability densities (see Materials and Methods) of the burst onsets (Figure 4A) and burst offsets (Figures 4B) phases. The values of probability density above its average level in the locomotor cycle (indicated by horizontal dotted lines in Figures 4A,B) formed peaks. Some of them were large and exceeded the double of the average value (indicated by horizontal dashed lines in Figures 4A,B), while others did not reach it. Peaks of the burst onsets (Figure 4A) fell into 4 phase ranges (I-IV), while peaks for the offset probability (Figure 4B) fell into 3 phase ranges (I, II and IV). These results are qualitatively similar to lower precision results of previous studies (Orlovsky & Feldman, 1972).

**FIGURE 4.**
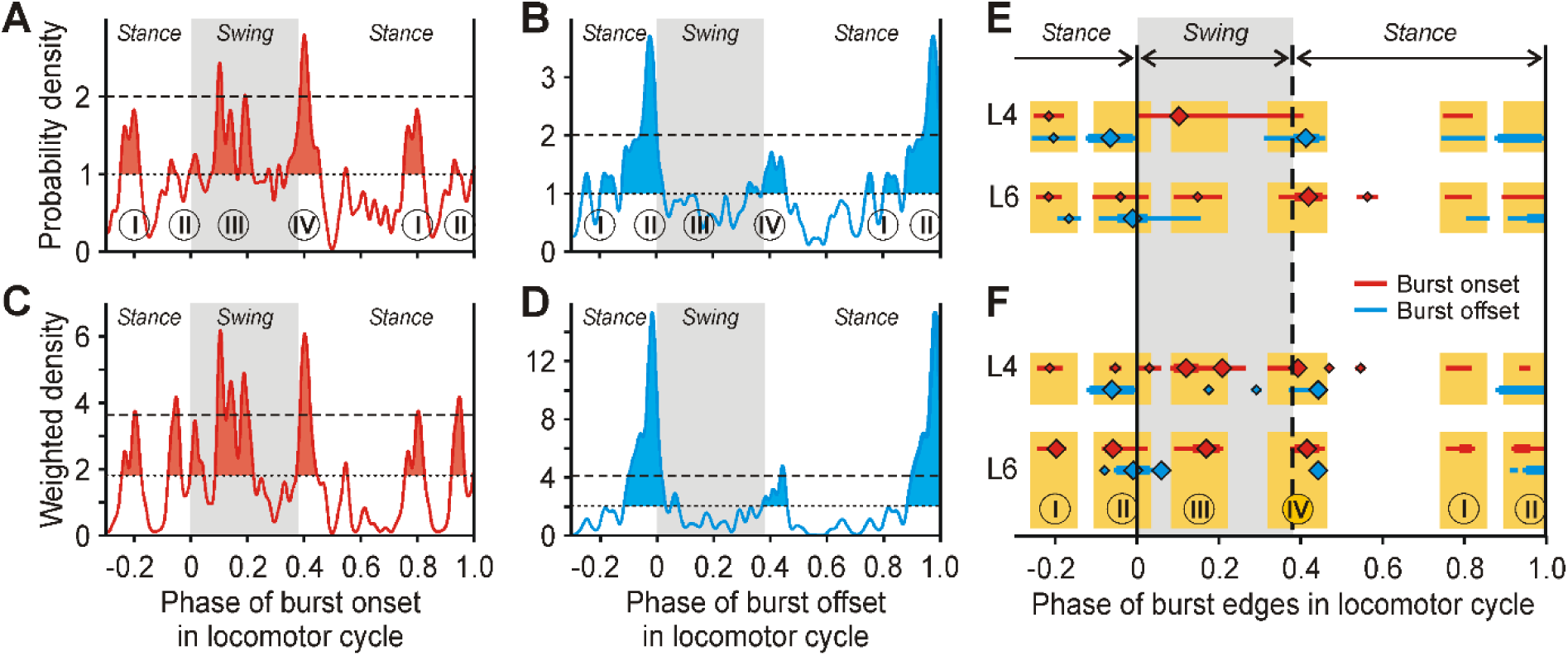
Important phases of the locomotor cycle. (A-D) Distributions of phase probability density (A,B) and phase weighted probability density (C,D) for the burst onsets (A,C) and offsets (B,D) for all (L4+L6) neurons. (E,F) Phases of peaks (parts of the curve above the average level indicated by dotted line in A-D) and major peaks (above the doubled average level indicated by dashed line in A-D) of probability (E) and weight (F) distributions are shown separately for L4 and L6 neurons. Thin and thick lines, and small and large diamonds indicate peaks and major peaks, and the peak tops, respectively. Only significant peaks were taken into account: if a peak would disappear after the exclusion of one neuron with *w* = 1, this peak was considered not significant. Additionally, two neighboring peaks were fused into one if they were separated by an insignificant trough that would disappear after the addition of one neuron with *w* = 1. Practically all peaks fell into 4 phase ranges I-IV (indicated by yellow in E,F).

The observed complex patterns of peaks can be explained either by real presence of many important phases in the locomotor cycle or by data noise. To reduce the noise, we examined the weighted phase distributions (see Materials and Methods), assigning larger weight to edges with lower *SD.* Thus, the weight of 1 was assigned to edges with *SD =* 0.1, which, on one hand, was the median value of both the onset and offset phases in our database, and on the other hand constituted a rather small part of the locomotor cycle (approximately one quarter of the swing or one sixth of the stance). The weight was higher than 9 (although never higher than 10) for edges with *SD* ≤ 0.01, which is a negligibly small variance, comparable to the instrument error in our experiments. Finally, the weight was lower than 0.3 (though always positive) for edges with *SD ≥* 0.2, which is a rather large variance (e.g., larger than a half of the swing).

In such weight distribution, data from approximately 30% of the entire population (neurons with large *SD*, which most likely were not critical for the locomotor pattern generation) was effectively filtered out. On the other hand, data from approximately 20% of the population (neurons with small *SD*, most likely, comprising the core of SLN) contributed more to the result. The weight distribution (Figures 4C,D) had the same peak locations as the probability distribution (Figures 4A,B) but the peaks were much clearer. Thus, fuzziness of the original edge distribution was most likely due to random data noise from neurons with unstable modulation (large *SD*).

In both L4 neurons and L6 neurons, most of the observed peaks of the onsets and offsets fell into the same 4 phase ranges (I-IV) (Figures 4E, F): range I with phases close to 0.8 part of the locomotor cycle, range II with phases close to 0 (beginning of the cycle), range III with phases close to 0.15 part of the cycle, and range IV with phases close to 0.4 part of the cycle. Thus, neurons were activated and inactivated not randomly in the locomotor cycle, but preferably within one of the 4 phase ranges. These 4 ranges were presumably related to discrete behavioral events: preparation for the limb lift-off (range I), the limb lift-off (range II), transition from the limb flexion to limb extension during swing (range III), and the limb touch-down (range IV).

### 3.3 | Classification of neurons based on their burst phases

To reveal groups of neurons with similar phase of activity modulation in the locomotor cycle, we considered only neurons with stable modulation that are most likely critically important for generation of locomotion. The following criteria for stability of modulation formulated in our previous study (Musienko et al., 2022) were used: (i) *SD* of the burst onset and/or burst offset should not be more than 0.1 part of the locomotor cycle. We accepted stability of only one edge of the burst, as a subset of the recorded neurons had ramp-up or ramp-down burst shape with stable burst offset and burst onset phase, respectively. (ii) The average coefficient of correlation between the profiles of the activity in individual locomotor cycles and in the entire activity histogram had to be no less than 0.6. According to these criteria, 126 out of 193 modulated neurons (65%) had stable modulation (“stable neurons”, Table 1). To the population of stable neurons, we applied a novel method of clustering that relies more on data with higher values of the phase weight (see Materials and Methods). As a result, activities of almost all stable neurons (91%, 115 out of 126 stable neurons) were classified into 19 *Clusters* (Table 2; Figure 5; Supplementary Figure 2A). In 11 neurons, activity phases were unique. We termed them “*Cluster* 0” and did not consider in the further analysis.

**FIGURE 5.**
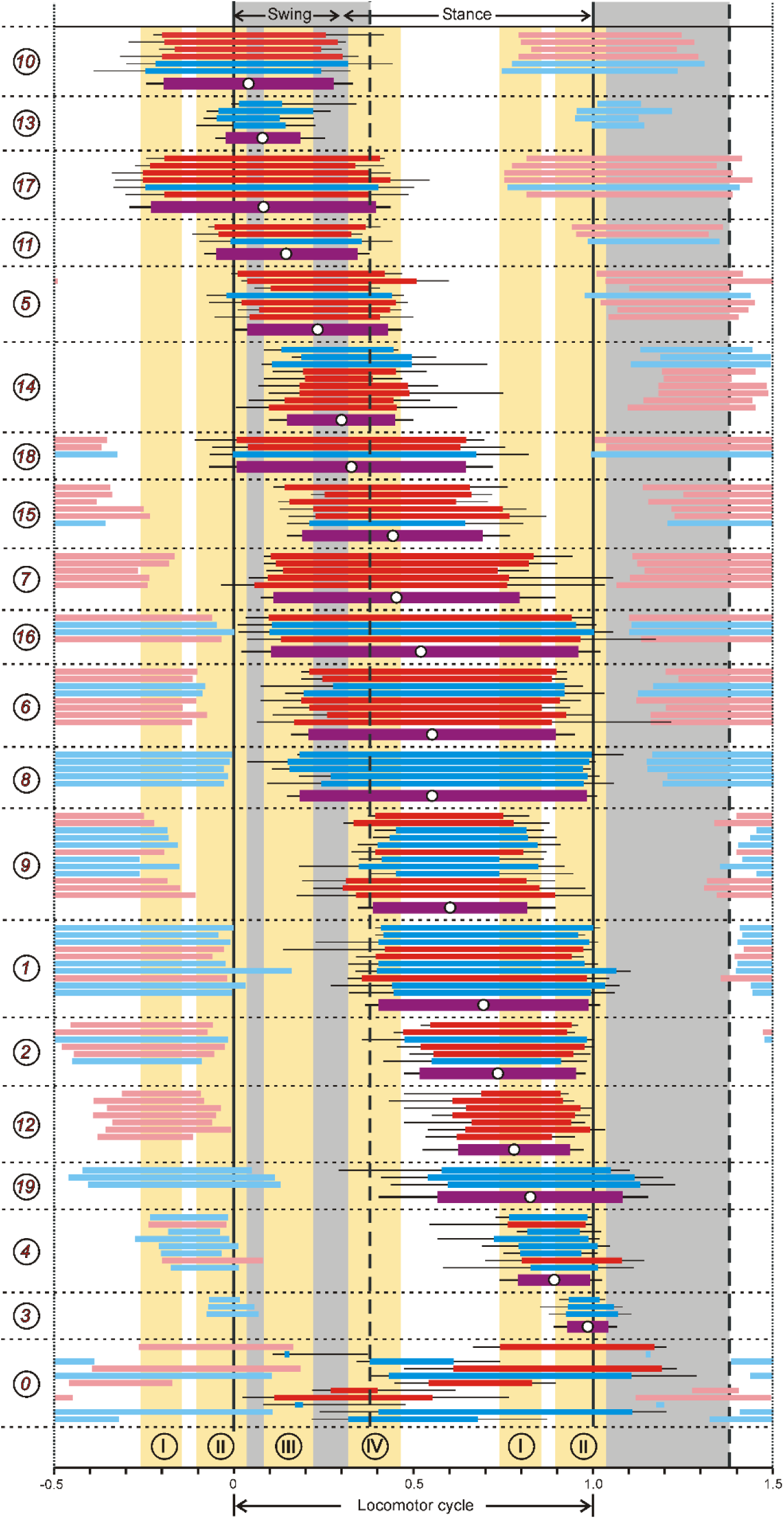
Phase distribution of sorted into *Clusters* bursts of individual neurons with stable modulation recorded in L4 and L6 segments in the normalized locomotor cycle. The mean bursts of individual *Clusters* with the middle point indicated (white circles) are shown by thick purple lines. Neurons of different *Clusters* are demarcated by dotted lines. *Clusters* are ordered according to the midburst phase. Numbers in circles on the left are *Cluster* numbers. Four phase ranges I-IV are indicated by yellow. Other abbreviations as in Figure 3.

**TABLE 2.** Rostro-caudal distribution of neurons belonging to different *Clusters*. Rostro-caudal distribution of neurons belonging to different *Clusters* as well as assorted neurons (*Cluster* 0), was compared to that of the entire recorded population using Fisher’s exact test (df = 1). Statistically significant (with *P* < 0.05) and visible but statistically insignificant (with 0.05 < *P* < 0.3, due to low numbers of neurons) biases in the neurons’ location are indicated, respectively, by bold and underlined *P* values. *n*, number of neurons.

| *Cluster #* | L4 (n) | L6 (n) | *P* |
| --- | --- | --- | --- |
| 1 | 3 | 7 | <u>0.19</u> |
| 2 | 4 | 2 | 0.69 |
| 3 | 0 | 3 | <u>0.09</u> |
| 4 | 2 | 6 | <u>0.14</u> |
| 5 | 6 | 1 | <u>0.23</u> |
| 6 | 6 | 2 | 0.47 |
| 7 | 5 | 0 | <u>0.07</u> |
| 8 | 0 | 5 | **0.02** |
| 9 | 6 | 6 | 0.77 |
| 10 | 4 | 2 | 0.69 |
| 11 | 2 | 1 | 0.99 |
| 12 | 7 | 0 | **0.02** |
| 13 | 0 | 4 | **0.04** |
| 14 | 6 | 3 | 0.73 |
| 15 | 5 | 1 | <u>0.23</u> |
| 16 | 2 | 2 | 0.99 |
| 17 | 5 | 1 | <u>0.23</u> |
| 18 | 2 | 1 | 0.99 |
| 19 | 0 | 3 | <u>0.09</u> |
| 0 | 5 | 6 | 0.54 |
| Total | 70 | 56 |  |

Since the neurons within each *Cluster* had similar activity phases, in the further analysis, we considered each *Cluster* as a functionally homogeneous set of neurons. For each *Cluster*, the phase of activity of the *Cluster* was characterized by weighted average phases of the burst onsets and offsets of neurons comprising the *Cluster* with the corresponding *SD*s (see Materials and Methods). The interval between these average phases was termed the “burst” of the *Cluster* (thick purple lines in Figure 5). As seen in Figure 5, the majority of *Clusters* had the burst onset in one phase of the locomotor cycle (swing or stance) and the burst offset in another phase. Only two *Clusters* (*2* and *12*) had bursts within the stance and no one within the swing.

### 3.4 Functional classification of *Clusters*

In different SLN models (Ijspeert, 2008) the locomotor rhythm is generated by neuronal classes with reciprocal inhibition, which are active out of phase. To reveal such pairs of *Clusters* active out of phase, for each *Cluster* we calculated its midburst phase (see Materials and Methods; white circles on thick purple lines in Figure 5). Then, for each pair of *Clusters*, we calculated the phase shifts between their midburst phases (see Materials and Methods as well as Supplementary Data pages 10-11 for specific details). Figure 6A shows the heatmap of the phase shift between midbursts in different *Cluster* pairs. All *Clusters* can be combined into 4 pairs of *Subgroups* active approximately out of phase (the corresponding phase shifts outlined by 4 dashed rectangles in Figure 6A): *Subgroup 1* (*Clusters 10, 13, 17*) is out of phase with *Subgroup 2* (*Clusters 16, 6, 8, 9*), *Subgroup 3* (*Clusters 11, 5*) – with *Subgroup 4* (*Clusters 1, 2*), *Subgroup 5* (*Clusters 14, 18*) - with *Subgroup 6* (*Clusters 12, 19*), and *Subgroup 7* (*Clusters 15, 7*) – with *Subgroup 8* (*Clusters 4, 3*).

**FIGURE 6.**
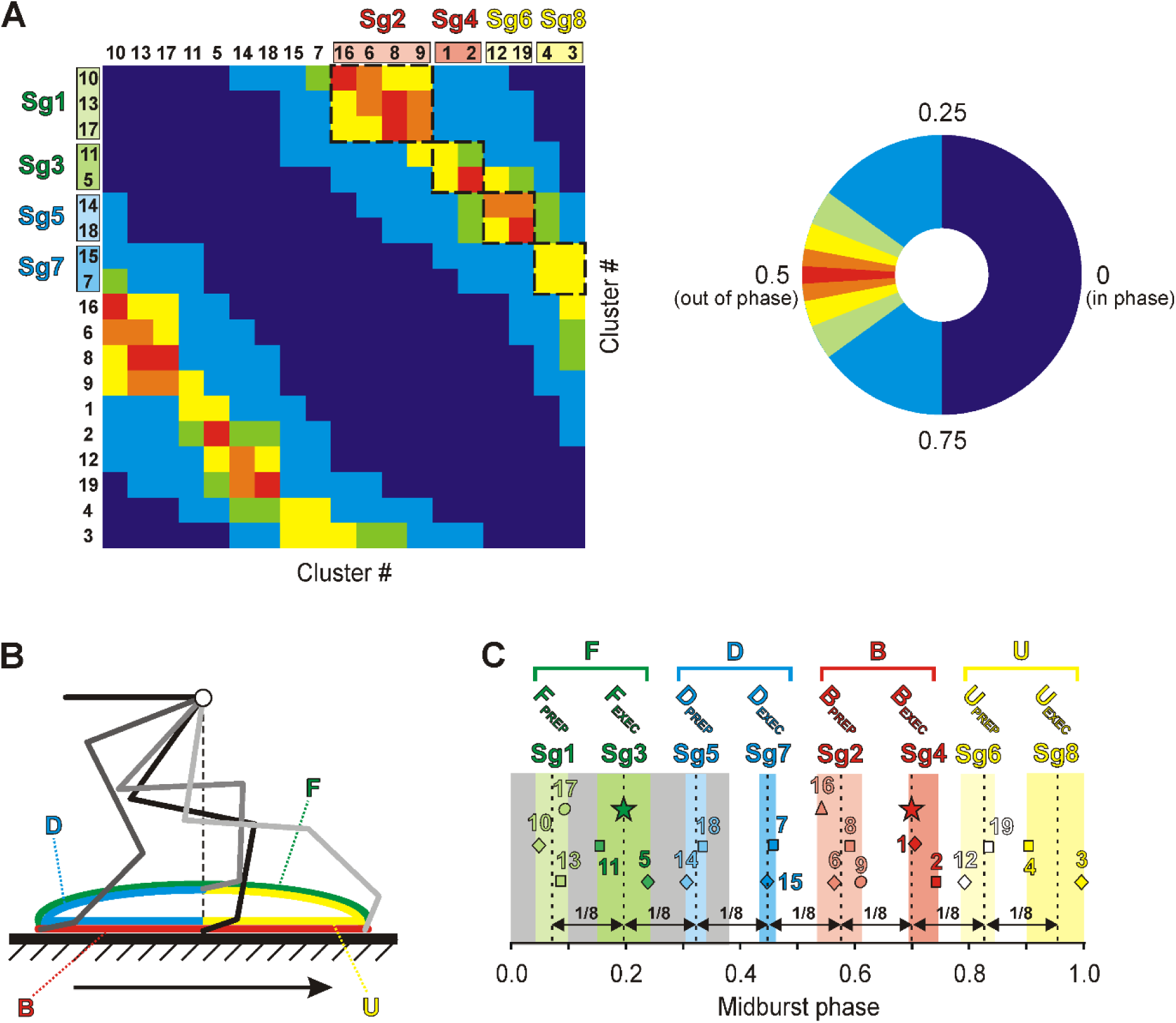
Functional classification of *Clusters*. (A) Heatmap of the phase shift between midbursts of different *Cluster* pairs. All *Clusters* can be combined into 4 pairs of *Subgroups* active approximately out of phase (with phase shifts close to 0.5, outlined by 4 dashed rectangles): Sgr1/Sgr2, Sgr3/Sgr4, Sgr5/Sgr6, Sgr7/Sgr8. (B) The paw trajectory during the locomotor cycle. Green, red, yellow and blue parts of the trajectory are, respectively, forward (F), backward (B), upward (U), and downward (D) limb movements. HC and VC, the horizontal and vertical components of the step. (C) *Subgroups* are formed by *Clusters* with similar midburst phases (each color band corresponds to one *Subgroup*). These *Subgroups* can be assigned functional roles (*F_PREP_, F_EXEC_, D_PREP_, D_EXEC_, B_PREP_, B_EXEC_, U_PREP_, U_EXEC_*), and they form four *Groups* (*F, D, B, U*) (see explanations in the text and Supplementary Figure 7). Green and red stars show the middle of the swing and stance, respectively.

*Clusters* constituting one *Subgroup* had similar midburst phases (Figure 6C). Phase ranges of midbursts of *Clusters* forming individual *Subgroups* (indicated by colored bands in Figure 6C) were shifted in relation to one another by approximately 1/8 of the locomotor cycle. To assign functional roles to the *Subgroups*, one can take into account the paw trajectory during the locomotor cycle (Figure 6B; Supplementary Figure 7). It can be divided into forward (the green part of the trajectory, F) and backward (the red part of the trajectory, B) movements corresponding to the horizontal component of the step (HC) or into two parts corresponding to the vertical component of the step (VC) that includes the movements of the limb when it is subjected to unloading and upward transition (the yellow part of the trajectory, U) and to downward transition and loading (the blue part of the trajectory, D), respectively. Phases of HC and VC are shifted by approximately 1/4 of a cycle (Figure 6B, Supplementary Figure 7). The existence of separate neural mechanisms generation VC and HC of the step was demonstrated earlier (Musienko et al., 2012). Thus, *Subgroups 3 and 4,* with activity phase ranges close to the middle of the swing and to the middle of stance, respectively, can be responsible for execution of the forward and backward paw movements, and correspondingly termed *F_EXEC_* and *B_EXEC_* (Figure 6C). *Subgroup 7,* with its activity following *F_EXEC_* by 1/4 and leading *B_EXEC_* by 1/4, executed the downward movement (limb extension during the swing second half, touchdown, and weight support during the stance first half), while *Subgroup 8* executed the upward movement (limb unloading during the stance second half, lift-off, and limb flexion during the swing first half) (Figure 6C). Correspondingly, *Subgroups 7* and *8* were termed *D_EXEC_* and *U_EXEC_.* The remaining *Subgroups 1, 5, 2,* and *6,* leading the corresponding executive *Subgroups* (*F_EXEC_, D_EXEC_, B_EXEC_, and U_EXEC_*) by 1/8 of the locomotor cycle (Figure 6C), can be assigned preparatory roles for the forward, downward, backward, and upward movements, respectively. Correspondingly, they were termed *F_PREP_, D_PREP_, B_PREP_,* and *U_PREP_.* Couples of the preparatory and executive *Subgroups,* presumably controlling movement in a specific direction (forward, backward, upward, downward) during the step cycle, form *Groups F, B, U,* and *D*, respectively (Figures 6C, 9A). *Groups F* and *B* comprise a network generating the HC of movement, while the VC network consists of *Groups D* and *U*.

Although in general, neurons forming different *Clusters* were intermingled in the gray matter, we found some specificity in their spatial distribution. Thus, neurons of *Clusters 7* and *12*, were found only in the spinal segment L4, while neurons of *Clusters 3, 8, 13* and *19* were found only in L6 (Figures 7 and 5, Table 2). In both L4 and L6, the majority of neurons forming the VC network (*Groups U* and *D*) were located in the intermediate zone of the gray matter, while the majority of neurons forming the HC network (*Groups F* and *B*) were located in the dorsal and ventral zones (Figure 7, Table 3). In both segments these differences in distribution of VC and HC neurons were statistically significant (Table 4).

**FIGURE 7.**
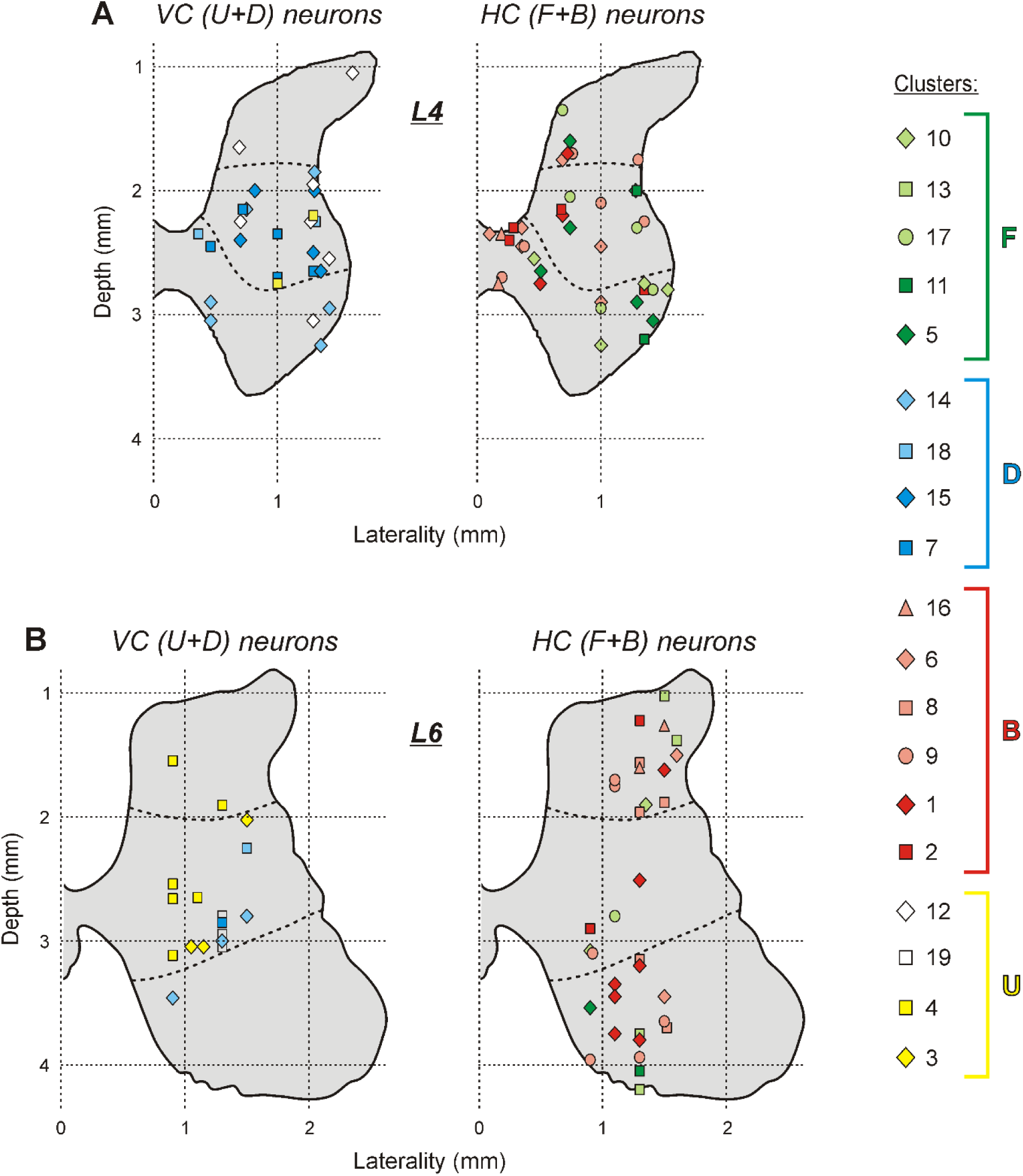
Location of individual neurons of different *Clusters* in the spinal cord. VC network neurons and HC network neurons are shown separately on cross-sections of the gray matter at spinal segments L4 and L6. Thick hatched lines demarcate the dorsal, intermediate, and ventral zones of the gray matter. Designations (shapes and colors) are as in Figures 6C.

**TABLE 3.**
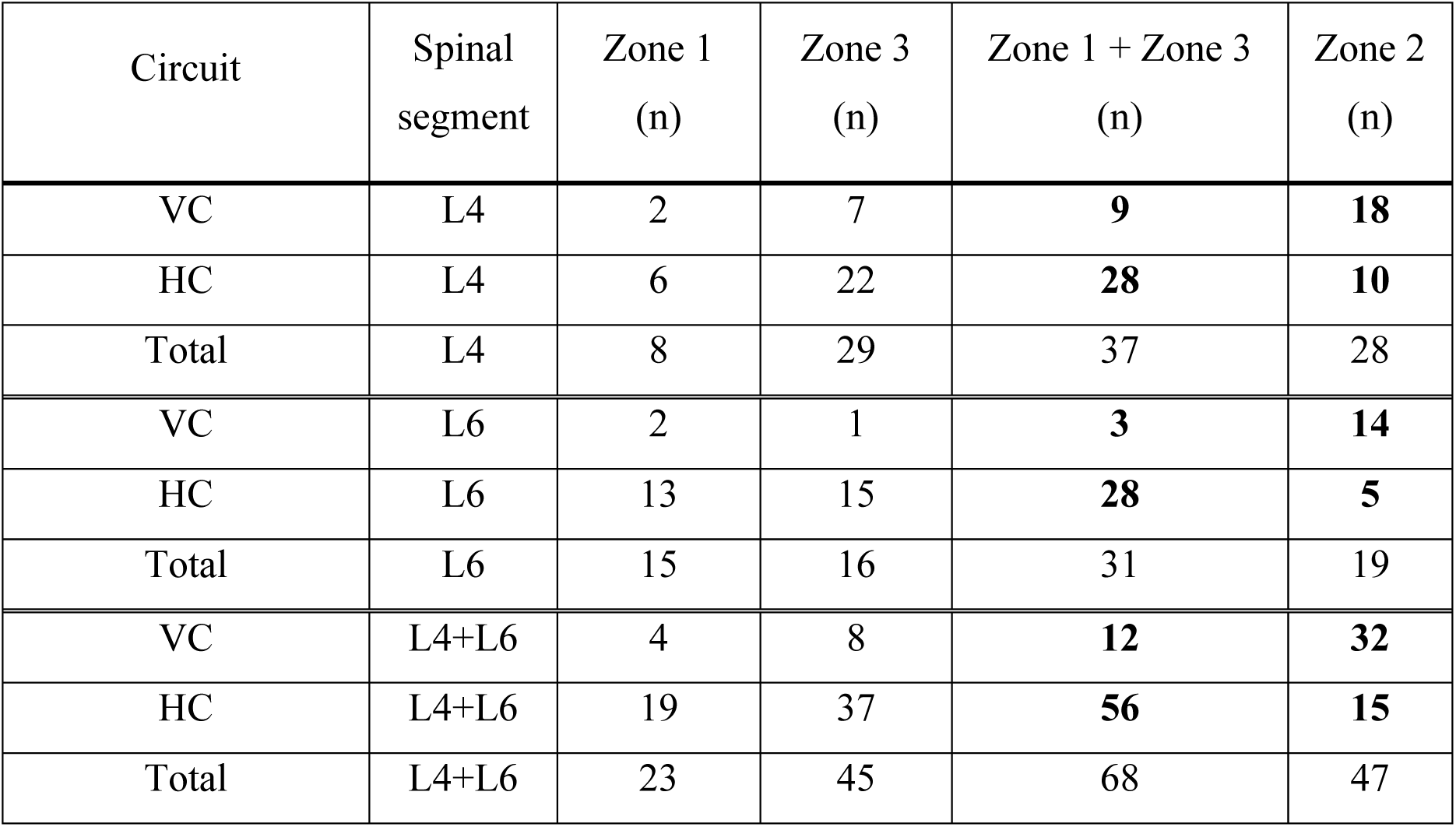
Rostro-caudal and dorso-ventral distribution of neurons comprising networks generating the vertical and horizontal components of stepping. VC and HC, networks generating, respectively, the vertical and horizontal components of stepping. The number of neurons (*n*) in each of three zones of the gray matter (dorsal, Zone 1; intermediate, Zone 2; ventral, Zone 3; Figure 8) is indicated. Numbers shown in bold are statistically compared in Table 4.

**TABLE 4.** Statistical comparison of the distribution of VC and HC neurons across the transverse section of the gray matter. The relative number of neurons in Zone 2 and Zones 1+3 (see Table 3) were statistically compared using Fisher’s exact test (df = 1). Values of *P* < 0.05 (statistically significant bias in the neurons’ location) are indicated by bold.

| Comparison | Spinal segment | *P* |
| --- | --- | --- |
| VC *vs* HC | L4 | **0.002** |
| VC *vs* HC | L6 | **0.0001** |
| VC *vs* HC | L4+L6 | **0.0001** |

### 3.5 Functional connections between *Clusters* and *Groups*

To reveal possible functional connections between *Clusters*, we analyzed short (less than 0.1 part of the cycle) phase shifts between the burst onsets/offsets in different *Cluster* pairs (Δ in Figures 8A,B). We suggested that one *Cluster* (*Pre* in Figure 8A) inhibited another *Cluster* (*Post* in Figure 8A) if the burst onset of the *Pre Cluster* was followed by the burst offset of the *Post Cluster* [activation of “presynaptic” inhibitory neurons inactivated the “postsynaptic” target; *Inactivation by inhibition* in Figure 8A(i)], or the burst offset of the *Pre Cluster* was followed by the burst onset of the *Post Cluster* [*Activation by disinhibition* in Figure 8A(ii)]. The latter can be explained by postinhibitory rebound that was demonstrated in spinal interneurons (e.g., Rivera-Arconada & Lopez-Garcia, 2015; Zhu et al., 2021). The functional connection was considered excitatory if the burst onset of the *Pre Cluster* was followed by the burst onset of the *Post Cluster* [*Activation by excitation* in Figure 8A(iii)] or if the burst offset of the *Pre Cluster* was followed by the burst offset of the *Post Cluster* [*Inactivation due to termination of excitation* in Figure 8A(iv)].

**FIGURE 8.**
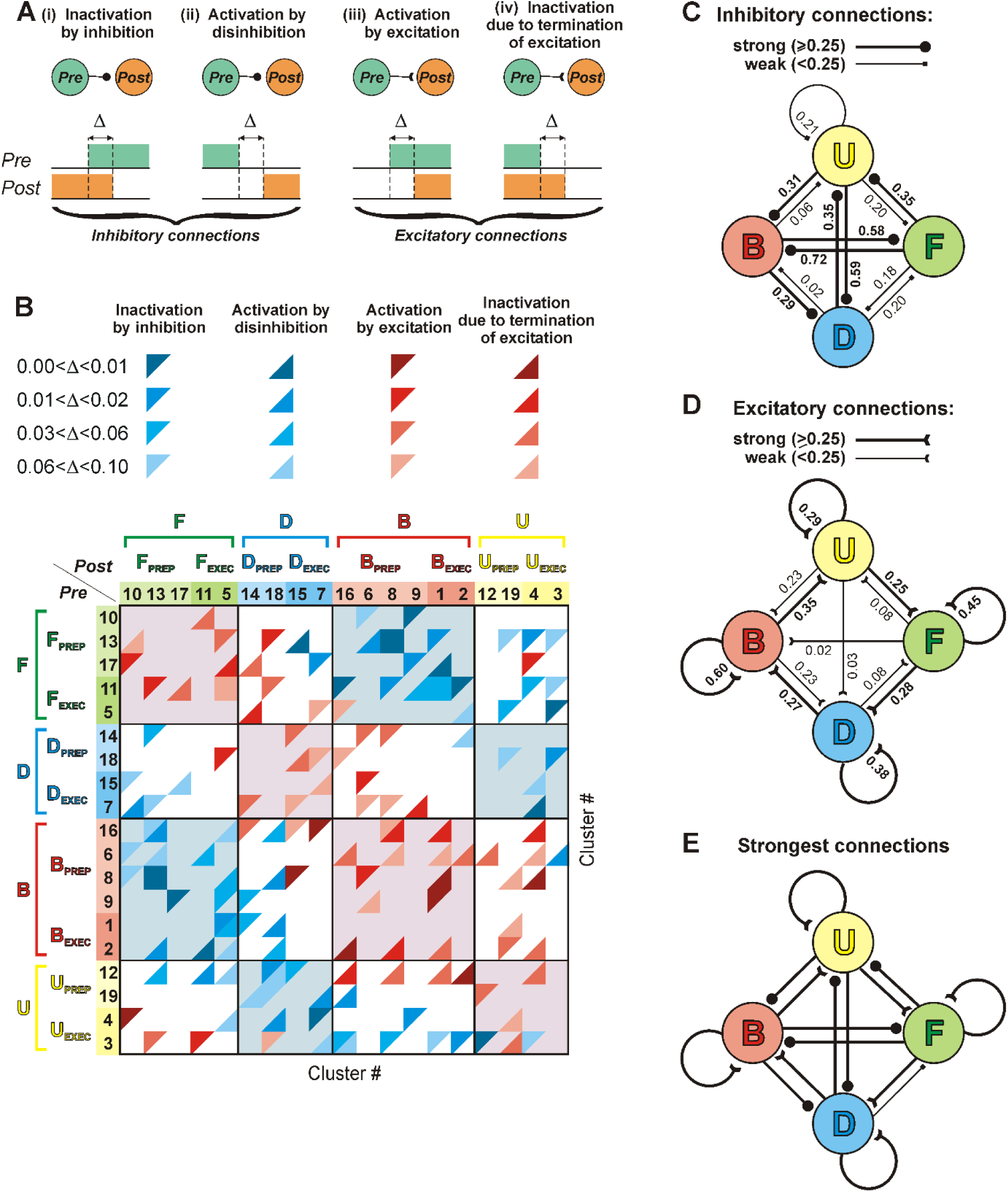
Functional connections between *Clusters*. (A) Revealing possible functional connections between *Clusters* (see Results for explanation). (B) Heatmap showing functional connections between different *Clusters*. Blue triangles, inhibitory connections; red triangles, excitatory connections. (C-E) A summary of the revealed inhibitory (C), excitatory (D), and strongest (E) functional connections between *Groups*. Numbers are the values of parameter *S* characterizing the strength of functional excitatory or inhibitory connections between the *Clusters* (see Results for details). *S* values exceeding the median level (0.25) correspond to “strong” connections (marked by bold), while connections with *S* below the median are considered “weak”.

Figure 8B shows the heatmap of revealed functional connections between different *Pre* and *Post Clusters* belonging to different *Groups. Cluster* pairs with all four types of abovementioned interactions (Figure 8A) were found. We did not observe any *Cluster* pair with both excitatory and inhibitory connections. Between *Pre* and *Post Clusters* of each *Group*, almost all revealed connections were excitatory (red triangles on the pink background in Figure 8B). Between *Pre Clusters* forming *Group U* and *Post Clusters* forming *Group D* (VC network), as well as between *Pre Clusters* forming *Group F* and *Post Clusters* forming *Group B* (HC network), almost all connections were inhibitory (blue triangles on the light blue background in Figure 8B), which likely explains the reciprocal activity within these pairs of *Groups*. Finally, between *Pre* and *Post Clusters* of *Groups* that belong to networks contributing to generation of different components of the step (e.g., between *Clusters* of *Group D* that belongs to VC and *Clusters* of *Group B* that belongs to HC) both inhibitory and excitatory connections were found (blue and red triangles on the white background in Figure 8B).

The values of the phase shifts between the burst onsets/offsets (reflecting the latencies of functional effects, Δ in Figure 8A,B) were different in different *Cluster* pairs. One can suppose that functional effects with shorter latencies are mediated by stronger connections, while those with longer latencies are mediated by weaker connections. To evaluate the strength of functional excitatory or inhibitory connections between the *Groups,* an integral parameter *S* (“strength”) was calculated: *S* = (4*N_4_* + 3*N_3_* + 2*N_2_* + *N_1_*) / *N*, where *N_4_*, *N_3_*, *N_2_*, and *N_1_* were the numbers of revealed connections with the latency in the ranges 0.00-0.01, 0.01-0.03, 0.03-0.06, and 0.06-0.10 parts of the cycle, respectively, and *N* was the total number of Δ measurements performed for a pair of *Groups*. The *S* values varied from 0.03 to 0.46 for excitatory connections and from 0.02 to 0.72 for inhibitory ones (Figure 8C,D). However, the median *S* values for both types of connections were almost the same (approximately 0.25). Connections between *Groups* with *S* value below and above 0.25 were considered as weak and strong connections, respectively (indicated by thin and thick lines, respectively, in Figure 8C,D).

Mutual inhibitory connections between *Groups* of VC (*U* and *D*) and between *Groups* of HC (*F* and *B*), as well as mutual excitatory connections between *Clusters* within each of *Groups,* were strong (Figure 8C,D). By contrast, excitatory and inhibitory connections between the majority of VC and HC *Groups* were anisotropic, e.g., *Group U* strongly excited and weakly inhibited *Group F*; by contrast, *Group F* strongly inhibited and weakly excited *Group U* (Figure 8C,D). Thus, one can suggest that *Groups* form a generating ring -*U-F-D-B*-(Figure 8E), in which each *Group* inactivates the preceding one and activates the next one.

So, although each *Cluster* of a *Group* is not homogeneous and contains neurons producing excitatory as well as neurons producing inhibitory effects, the functional effect of one *Group* upon another *Group* is determined by the strongest excitatory and inhibitory connections. Thus, activation of *Group* U leads to activation of all its neurons (due to mutual excitatory connections) leading to inhibition of *Groups* B and D that results in lifting of the limb above the ground. Simultaneously, activated *Group* U activates *Group* F. Activation of *Group* F leads to activation of all its neurons (due to their mutual excitatory connections) that leads to inhibition of *Group* U and *Group* B and resulting in forward progression of the limb. Simultaneously, *Group* F activates *Group* D that in turn inhibits *Group* U and *Group* F that results in downward movement of the limb. Simultaneously, *Group* D activates *Group* B that in turn inhibits *Groups* D and F resulting in the backward limb movement. Activated *Group* B simultaneously activates *Group* U and so on.

## 4 Discussion

### 4.1 Neurons comprising the spinal locomotor network (SLN)

In the present study we analyzed the spinal locomotor network (SLN) generating real forward locomotion with close to normal movement-related sensory feedback. Spinal interneurons form the core of this network. It should be noted that we discriminate between SLN and the central pattern generator (CPG, the part of the spinal network capable of generating the locomotor pattern without sensory feedback). SLN includes CPG but is not limited to CPG.

There is some discrepancy in results of recording of spinal interneurons with activity modulated in the locomotor rhythm. Studies in which spinal interneurons were recorded during forward locomotor movements [e.g., earlier studies in which a small database was obtained during real forward locomotion in cats with intact sensory feedback and after deafferentation (Orlovsky & Feldman, 1972), studies on spinal cats performing forward air-stepping (AuYong et al., 2011), as well as the present study] demonstrated rather uniform distribution of phases of interneuronal activity across the locomotor cycle. By contrast, recording of spinal interneurons during spontaneous episodes of fictive locomotion in rabbits (Viala et al., 1991) and cats (Baev et al., 1979) revealed two clusters of neurons with activity phases almost coincident with either flexor or extensor motoneuronal activity. Unfortunately, in the last two studies only motor nerves to flexor and extensor muscles of the ankle joint were recorded that did not allow to determine the type of generated locomotor pattern (i.e. whether the motor pattern corresponded to forward stepping, backward stepping, or in place stepping). One of possible explanations of the discrepancy between the data obtained during fictive locomotion and forward locomotion with intact movement-related sensory feedback, is that in immobilized animals, neurons were recorded during in place locomotion characterized by simple reciprocal activity of flexor and extensor muscles around all joints of the limb (Lyakhovetskii et al., 2021). Earlier we demonstrated that spinal networks generating the forward and backward stepping differ to some extent (Merkulyeva et al., 2018; Musienko et al., 2022).

In our study, we focused on interneurons with stable modulation of activity that most likely represent the essential elements of SLN. We excluded neurons with sparse firing from our analysis. It has been suggested that such neurons may play an important role in the rhythm generation (Strohmer et al., 2024). Unfortunately, adequate analysis of sparse activity would require recording of much longer episodes than used in the present study. This can be a goal of future studies.

Since spinal interneurons were recorded during real forward locomotion in the presence of normal movement-related sensory feedback, we cannot exclude that locomotion-related modulation of activity in a part of these neurons was driven by the sensory feedback. There are numerous findings (e.g., Grillner & Rossignol. 1978; Whelan et al., 1995; Hiebert et al., 1996) strongly suggesting that in real stepping, the timing of different events in the step cycle is determined not by CPG activity (Orlovsky et al., 1999) but rather by afferent signals about the limb movement. Thus, sensory driven neurons represent an important part of SLN generating forward locomotor movements. However, the majority of interneurons recorded in our study were similar to those revealed during locomotion in deafferented cats (Orlovsky & Feldman, 1972; Dai et al., 2005) in terms of location (the ventral and intermediate zones of the gray matter, Figure 7), and in terms of distribution of the activity phases across the locomotor cycle. Therefore, one can suppose that most of our SLN neurons belonged to CPG.

We correlated activity of recorded neurons with the locomotor cycle of the ipsilateral limb. We cannot exclude that a part of recorded neurons were commissural interneurons involved in control of the contralateral limb. However, the majority of neurons were recorded outside of areas in which commissural interneurons are located. Thus, we do not think that a few commissural neurons in our database can affect the interpretation of our results.

### 4.2 Possible functional roles of neuronal *Clusters*

Our analysis revealed 4 phase ranges in the cycle (Figure 4) when many neurons were turned on and off. Onset and offset phases of the majority of neuronal *Clusters* revealed in the present study fell into the abovementioned 4 phase ranges (Figure 5). Thus, activity of SLN neurons is changed mainly around four critical moments in the locomotor cycle that demarcate F, E1, E2, and E3 phases of the cycle revealed by kinematic analysis (Engberg & Lundberg, 1969). Most likely, changes in neuronal activity within range I contribute to transition from weight support (E2 phase) to limb unloading (E3 phase) as well as to preparation for the following limb lift-off. Changes in activity within range II contribute to transition from the stance (E3 phase) to the swing phase (F phase; the limb lift-off) as well as to preparation for the following limb forward transfer. Activity changes within range III may contribute to transition from the limb flexion (F phase) to the knee and ankle extension during swing (E1 phase) as well as to preparation for the following touch-down and weight acceptance. Finally, activity changes within range IV may contribute to transition from the swing (E1 phase) to the stance phase (E2 phase; the limb touch-down). In contrast, transition from limb shortening to limb elongation during stance (in the midstance) is less precisely controlled: neurons of *Clusters 19, 12* and *18, 15* that are turned, respectively, on and off in this part of the cycle (Figure 5) are relatively not numerous, and their on- and offset phases are defined not very precisely (note the larger *SD*s in these *Clusters* in Figure 5).

Analysis of the motor pattern of forward stepping revealed a few (2-5 in different studies) motor modules or muscle synergies that underly forward locomotor movement (Dominici et al., 2011; Ting et al., 2015; Aoi et al., 2013). It was suggested that their phases represent important events in the locomotor cycle. While there is a consensus that this structure exists in motor patterns, however, how it arises, whether it reflects neuronal structure, and whether it is functionally relevant are sources of lively debate.

Suggested by our analysis functional effects of each *Cluster* can be excitatory on some *Clusters* but inhibitory on others (Figure 8B). Thus, one can assume that each *Cluster* contains several types of neurons differing in projections as well as neurotransmitters.

The activity phase of some *Clusters* (Figure 5) coincided with the activity phase of specific limb muscles (Krouchev et al., 2006; Markin et al., 2012). For example, activity of *Cluster* 3 coincided with activity of muscles initiating the swing (cluster 2 in Figure 7, Krouchev et al., 2006, and cluster 5 in Figure 5B, Markin et al., 2012) and *Cluster 14* was active at the end of swing and the beginning of stance, similar to muscle cluster 5 in Figure 7, Krouchev et al., 2006, and cluster 7 in Figure 5B, Markin et al., 2012. Neurons of such *Clusters* could be the last order premotor interneurons. However, the other *Clusters* (e.g., *7, 10, 17*) had no motoneuronal counterparts, and thus, they probably do not directly affect the motoneurons.

### 4.3 Conceptual model of SLN

Our previous study, in which SLN was activated by epidural stimulation, demonstrated the existence of separate neural mechanisms generating vertical (VC) and horizontal (HC) components of a step (Musienko et al., 2012). The present study suggests that according to their activity phases, all interneuronal *Clusters* can be assigned either to VC or HC parts of SLN (Figure 9A). Moreover, the VC consists of a pair of reciprocally inhibiting *Group U* (up) and *Group D* (down) (Figure 9B), while the HC consists of a pair of reciprocally inhibiting *Group F* (forward) and *Group B* (backward) (Figure 9C). Thus, in contrast to previous studies (Orlovsky & Feldman, 1972; Baev et al., 1979; Viala et al., 1991; AuYong et al., 2011), we found several pairs of reciprocally inhibiting neuronal *Groups* with activity in different and specific parts of the step cycle.

**FIGURE 9.**
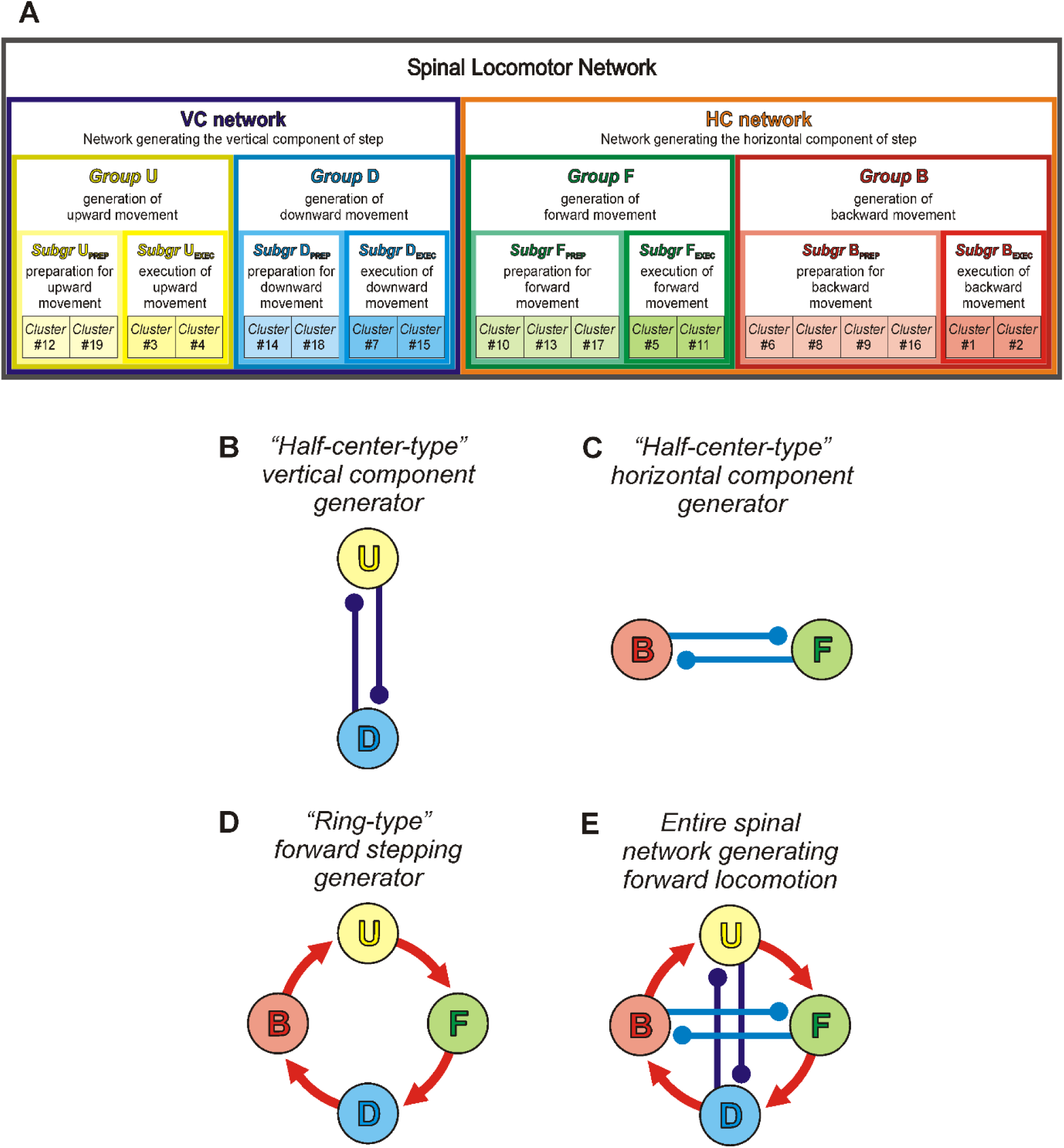
Summary of obtained results. (A) Hierarchical organization of the spinal locomotor network (SLN). (B-E) A hypothetical model of SLN based on revealed connections between *Clusters*. (B) A part of SLN generating the vertical component of stepping. The up (U) and down (D) half-centers are active out of phase (the sequence of activity: up – down – up – down – …) due to reciprocal inhibition. (C) A part of SLN generating the horizontal component of stepping. The forward (F) and backward (B) half-centers are active out of phase (the sequence of activity forward – backward – forward – backward – …) due to reciprocal inhibition. (D) Due to anisotropic inhibition and excitation in pairs U/F, F/D, D/B, and B/U, activity is transferred in ring manner (red arrows show the sequence of activity up – forward – down – backward – up – …). (E) The entire SLN for forward locomotion combines these two half-center-type networks and one ring-type network. Color coding of *Clusters, Subgroups* and *Groups* is the same as in Figure 6C.

In addition, anisotropic excitatory and inhibitory connections (Figures 8C,D) result in formation of the *-U-F-D-B-* generating ring (Figure 9D). For example, when *Group U* is active, it will inactivate *Groups D* and *B* through the strong inhibitory connection and activate *Group F* through the strong excitatory connection (Figure 8E). Similarly, in its turn, activation of *Group F* will result in inactivation of *Groups U* and *B*, and activation of *Group D* (note that at this phase of the cycle, due to inactivation of *Group U*, *Group D* is no longer inhibited by *Group U*). Thus, activity will be conveyed from *U* to *F,* then from *F* to *D*, and so on (red arrows in Figure 9D). Correspondingly, transition of the neuronal activity from *Group U* to *Group F* will result in transition from generation of F phase (limb lift off) to generation of E1 phase (forward protraction of the limb). Transition of activity from *Group F* to *Group D* and then to *Group B* will lead to transition from generation of E1 phase to generation of E2 phase (the limb tough down, loading and beginning of the backward movement). Finally, transition of activity from *Group B* to *Group F* will lead to transition from generation of E2 phase to generation of E3 phase and then F phase (the limb unloading, end of the backward movement, and the limb lift-off, respectively).

It was demonstrated that the network generating the VC of step (*Groups U* and *D*) can be activated selectively that leads to in place stepping (Musienko et al., 2012). However, it is not clear whether only one or both VC *Groups* or only some *Clusters* of the *Groups* are rhythmogenic, while the others follow their rhythm, or if the rhythm is an emergent property of a part of or the entire VC network. Also, it is not clear whether the HC network (*Groups F* and *B*) alone can generate the rhythmic horizontal component of steps or if the HC network is driven by the VC network through the ring connections. These are questions for future studies.

It is interesting to note that almost all connections in the ring are not unidirectional but just anisotropic. One can expect that changing the efficacy of these connections (strengthening the weaker while weakening the stronger ones) will reverse the direction of the activity propagation (red arrows in Figure 9D will be reversed), resulting in a change of the direction of locomotion from forward to backward.

We also found that spinal interneurons forming the network generating VC and those forming the network generating HC of the step have significantly different distribution across the gray matter of the spinal cord. Most VC neurons (*Groups U* and *D*) were located in the intermediate part of the gray matter while the majority of HC neurons (*Groups F* and *B*) – in the dorsal and ventral parts of the grey matter (Figure 7).

There are a number of models explaining the generation of locomotor movements, which are debated. Some earlier hypothetical models suggested that the locomotor rhythm can be generated by three or more neuronal elements forming a closed ring (Kling & Székely, 1968). Later, a model was suggested, in which the neuronal activity is continuously cycling through all phases of locomotor cycle (Linden et al., 2022). The most popular model based on numerous studies of locomotor networks in animal models of different complexity, suggests that a rhythm-generating part of the locomotor network consists of just two populations (half-centers) of neurons with mutual inhibitory connections (Brown, 1914; Jankowska et al., 1967; Lundberg, 1981; Ijspeert, 2008). Experimental observations demonstrated that the original half-center hypothesis in its simplest form cannot account for specific profiles of EMG activity (Perret & Cabelguen, 1980) and non-resetting deletions of activity of specific muscles (Lafreniere-Roula & McCrea, 2005). Therefore, a more advanced SLN model (Rybak et al., 2006; McCrea & Rybak, 2008; Rybak et al., 2015) includes a rhythm generator of the half-center type, and an output stage responsible for formation of the locomotor EMG pattern. Another SLN model (Grillner, 1985, 2006, 2021) suggests that the locomotor network contains a set of rhythm-generating units, each controlling muscles of a particular joint, and specific interactions between these units result in generation of a particular locomotor pattern. Finally, CPG controllers incorporating pattern formation models simulating modular muscle activations with multiple patterns loosely corresponding to different locomotor phases have been proposed as well (Aoi et al., 2013).

In the present study we suggested a new model of SLN. However, we think that our experimental data are compatible with all other models. Each of the revealed *Clusters* can be rhythmogenic and thus represent a unit generator or a part of a unit generator from Grillner’s model. We cannot exclude that just one *Cluster* or a pair of mutually inhibiting *Clusters* are rhythmogenic, while the other *Clusters* represent different parts of the pattern formation layer from Rybak’s model. It is also possible that the minimal rhythmogenic circuit must include *Clusters* from all 4 *Groups* to form a minimal rhythmogenic ring. Future studies will clarify these issues.

Recent advances in genetics led to initiation of numerous studies devoted to identification of components of the spinal locomotor network based on transcription factor expression (for review, Kiehn, 2016). It turned out that no single genetically identified type of spinal interneurons has been found to be solely responsible for the locomotor rhythm generation or control of the step direction (Rancic & Gosgnach, 2021). It was suggested that the locomotor rhythm is generated by a set of genetically heterogeneous populations (Kiehn, 2016; Dougherty et al., 2013; Caldeira et al., 2017). Thus, there is evidence that at least two genetically identified populations of interneurons, glutamatergic Hb9-expressing interneurons and Shox2 non-V2a interneurons (Shox2+ interneurons that do not co-express Ch10), may be involved in generation of the locomotor rhythm (Dougherty et al., 2013; Caldeira et al., 2017). By contrast, it was demonstrated that two genetically identified populations, V1 and V2b interneurons are required for the flexor-extensor alternation during locomotion (inhibition of flexors is mediated by V1 interneurons and of extensors by V2b interneurons) (Britz et al., 2015).

To conclude, in the present study, by using a novel method of analysis of neuronal activity recorded during real forward locomotion, we have revealed groups of spinal putative interneurons presumably responsible for generation of the vertical and horizontal components of a step, their possible functional connections, and their contribution to generation of limb movement in different parts of the locomotor cycle, as well as roughly demarcated their locations. On the basis of the obtained results, a novel conceptual model for generation of forward locomotion has been suggested. The obtained data can be used as a benchmark for computational models of locomotor neuronal networks. Developed in the present study method of analysis can be used for analysis of rhythmical activity of neurons related to other motor functions (e.g., scratching, paw shaking, etc.).

### CRediT AUTORSHIP CONTRIBUTION STATEMENT

**Pavel E. Musienko:** Conceptualization, Funding acquisition, Investigation, Methodology, Writing – review and editing. **Oleg V. Gorskii:** Investigation, Methodology, Writing – review and editing. **Tatiana G. Deliagina:** Conceptualization, Funding acquisition, Investigation, Methodology, Validation, Visualization, Writing – original draft. **Pavel V. Zelenin:** Conceptualization, Formal analysis, Investigation, Methodology, Validation, Visualization, Writing – original draft.

## ACKNOWLEDGEMENTS

This work was supported by the St. Petersburg State University, St. Petersburg, Russia (projects: 94030803, 104623591) to PEM; by grant from NIH (R01 NS-100928) to TGD and PEM; by grant from NIH (R01 NS-064964) to TGD; by grants from Swedish Research Council (2020-02502) to TGD; by the Russian Science Foundation grant 22-15-00092 to PEM; by Sirius University of Science and Technology project: NRB-RND-2115 to PEM. We thank Natalia Merkulyeva for helping with the histological evaluation.

## DATA AVAILABILITY STATEMENT

The data that support the findings of this study are available from the corresponding author upon reasonable request.

## Abbreviations

SLN: spinal locomotor network
MLR: mesencephalic locomotor region
CPG: central pattern generator
HC: horizontal component of the step
VC: vertical component of the step

