## Supplementary Data for "Functional organization of the spinal locomotor network based on analysis of interneuronal activity"

### Musienko et al., Supplementary Data

#### ***Contents:***

I. Supplementary description of Materials and Methods.

1. *Conversion of a time sequence of spikes into the instantaneous rate* …..………… 2
2. *Definition of the phase of a locomotor cycle* ……..……………………………….. 3
3. *Estimation of the burst and interburst periods within a cycle of locomotion* …..….. 4
4. *Calculation of the mean and the standard deviation (SD) for phases of the burst onset and offset for an individual neuron*..……………………………...... 5
5. *Conversion of the burst edge (on/offset) SD value into the weight* ……….......... 6
6. *Analysis of the population distribution of burst onset and offset phases* ……......... 7
7. *Cluster analysis* ……........................................................................................ 8
8. *Combining Clusters into Subgroups* ..................................................................... 10

II. Supplementary Figures.

Supplementary Figure 1. *Conversion of the edge (on/offset) SD value into the weight*………………………………………… ……… 12

Supplementary Figure 2. *Combining stable neurons into Clusters*…………………… 13

Supplementary Figure 3. *Technical details for clustering of stable neurons*……………………………….…………………… 14

Supplementary Figure 4. *Search for Clusters active in anti-phase*…………….…….... 15

Supplementary Figure 5. *Combining Clusters into Subgroups* ………………….…….. 16

Supplementary Figure 6. *Cost for different combinations of Clusters into Subgroups* …………………….………………… 17

Supplementary Figure 7. *Vertical and horizontal components of the locomotor movements* ………………….………………… 18

### Supplementary description of Materials and Methods

#### 1. Conversion of a time sequence of spikes into the instantaneous rate

A time sequence of $N$ spikes that occurred at time instants $t_i$ ($i = 1, 2, \ldots, N$) can be represented as a raw density $raw(t)$:

$$raw(t) = \sum_{i=1}^{N} \delta(t - t_i)$$

where $\delta(t)$ is the Dirac delta function.

The raw density was used to calculate the "step" instantaneous frequency $step(t)$:

$$step(t) = \sum_{i=1}^{N-1} \frac{H(t - t_i)H(t_{i+1} - t)}{t_{i+1} - t_i}$$

where $H(t)$ is the Heaviside step function. Then the final instantaneous rate $f(t)$ was obtained by sliding averaging with a 15 ms window, which is equivalent to convolving:

$$f(t) = \int step(t - s)k(s)ds$$

with a "rectangular" kernel $k(s)$:

$$k(s) = \frac{H(s + 0.015)H(0.015 - s)}{0.03}$$

We have routinely used this method since 2008, as it was computationally simple and still produced reasonable results. For regular spiking (with constant interspike intervals) it produces perfectly constant frequency. Some examples of real spiking activity and the corresponding instantaneous rate are presented in **Figure 2A,B**.

#### 2. Definition of the phase of a locomotor cycle

We normalized the entire cycle duration to unity. For the cycle duration, average ± *SD* was 0.89 ± 0.10 s, minimum 0.67 s, maximum 1.10 s. Variability of the cycle duration within an episode was 0.064 ± 0.032 s, minimum 0.020 s, maximum 0.130 s.

The swing was normalized to 0.38 of the entire cycle. For the swing proportion, average ± *SD* was 0.38 ± 0.05, minimum 0.28, maximum 0.43. Variability of the swing proportion within an episode was 0.053 ± 0.012, minimum 0.035, maximum 0.067.

This double normalization of the locomotor cycle improves the comparability of different steps: e.g., activity around the take-off moment in one cycle is compared with activity around the take-off moment in the other cycles; the same is true for other important moments in the locomotor cycles (the middle of the swing, the paw touch-down, the middle of the stance). On the other hand, such normalization produces minimal distortions of the firing profile.

Explicitly, the procedure of double normalization was the following. For an episode of locomotion that included $N$ steps we had 3 sets of absolute time points: starts of steps (for which we took the swing starts) $\{a_i\}$, starts of step stance (which are, obviously, equal to the swing ends) $\{b_i\}$, ends of steps (for which we took the stance ends) $\{c_i\}$ ($i$ = 1, 2, …, $N$). Obviously, $c_i = a_{i+1}$ for $i$ = 1, 2, …, $N$-1. Then for any time moment $t$ that belongs to the $i$'th locomotor cycle (that is $a_i \leq t \leq c_i$) its phase $\varphi$ was calculated with a formula:

$$\begin{cases} \varphi = 0.38\dfrac{t-a_i}{b_i-a_i}, & if\ a_i \leq t \leq b_i \\ \varphi = 0.62\dfrac{t-b_i}{c_i-b_i} + 0.38, & if\ b_i \leq t \leq c_i \end{cases}$$

For any phase $\varphi$ (which, by the definition above, was in the range from 0 to 1), we considered values $\varphi$, $\varphi$+1, $\varphi$-1, $\varphi$+2, $\varphi$-2, … as equivalent. This can be seen, for example, in **Figure 3.**

#### 3. Estimation of the neuronal burst and interburst periods within a cycle of locomotion

For each locomotor cycle the instantaneous rate depending on time $f(t)$ was converted into the instantaneous rate depending on the cycle phase $f(\varphi)$ (Figure 2B,C). For the phase $\varphi$ being in a range:

$$0 \le \varphi \le 1$$

Then we approximated $f(\varphi)$ with the best two-level rectangular fit $rect(\varphi)$:

$$rect(\varphi) = f_1 H(\varphi - \varphi_1) H(\varphi_2 - \varphi) + f_2\big(1 - H(\varphi - \varphi_1) H(\varphi_2 - \varphi)\big)$$

where $H(\varphi)$ is the Heaviside step function, and $f_1$, $f_2$, $\varphi_1$, $\varphi_2$ are constant parameters. We chose not to consider bursts or intebursts that occupy more than 97% of the cycle (phases $\varphi_1$, $\varphi_2$ had to be not very close to one another):

$$0.03 \le \varphi_2 - \varphi_1 \le 0.97$$

This choice was arbitrary but seemed reasonable. As the cycle duration was approximately 1 s so we did not considered parameters which determined bursts and interbursts shorter than 30 ms.

Parameters $f_1$, $f_2$, $\varphi_1$, $\varphi_2$ were chosen to minimize the cost function $C(f_1, f_2, \varphi_1, \varphi_2)$, namely the squared difference between the instantaneous rate $f(\varphi)$ and the approximation $rect(\varphi)$:

$$C(f_1, f_2, \varphi_1, \varphi_2) = \int_0^1 \big(f(\varphi) - rect(\varphi)\big)^2 d\varphi$$

The minimum and maximum of $f_1$ and $f_2$, were termed, correspondingly, the burst frequency $f_{\mathrm{BURST}}$ and the interburst frequency $f_{\mathrm{INTERBURST}}$:

$$f_{BUSRT} = \max(f_1, f_2)\,, f_{INTERBUSRT} = \min(f_1, f_2)$$

The burst onset phase $\varphi^{on}$ and the burst offset phase $\varphi^{off}$ were defined, correspondingly:

$$\varphi^{on} = \begin{cases} \varphi_1, f_1 > f_2 \\ \varphi_2, f_1 < f_2 \end{cases}, \varphi^{off} = \begin{cases} \varphi_2, f_1 > f_2 \\ \varphi_1, f_1 < f_2 \end{cases}$$

The phases $\varphi^{on}$ and $\varphi^{off}$ were estimated with the precision of $10^{-3}$.

#### 4. Calculation of the mean and the standard deviation (SD) for phases of the burst onset and offset for an individual neuron

For a set of phases (e.g., the burst onset phases, or the burst offset phases) $\{\varphi_k\}$ ($k = 1, 2, \ldots, N$) we calculated the mean $<\varphi>$ and the standard deviation *SD* using the circular statistics approach. Explicitly,

$$\langle\varphi\rangle = \frac{1}{2\pi}Arg(\bar{z}), \qquad SD = \frac{1}{2\pi}\sqrt{-2ln|\bar{z}|}, \qquad \text{where } \bar{z} = \frac{1}{N}\sum_{k=1}^{N} e^{2\pi i\varphi_k}$$

The minor difference in our formulas, compared to the standard formulas that are used in the circular statistics, was the scaling factor 2π: in traditional approach the phase varies in the range [0, 2π], while according to our definition the phase varied in the range [0, 1].

#### 5. Conversion of the burst edge (on/offset) SD value into the weight

In the present study, in addition to analysis used by the other authors, we propose one novelty: assigning more weight to the data with lower variability (smaller *SD*). With this purpose, for the mean phase $\varphi_i$ with $SD_i$, a new parameter, the weight, $w(SD_i)$, was calculated:

$$w(SD_i) = \frac{10}{1 + 900{SD_i}^2}$$

To understand the origin of this formula for the weight, one can transform it:

$$w(SD_i) = \frac{10}{1 + 900{SD_i}^2} = \frac{10}{1 + (30SD_i)^2} = \frac{10}{1 + 10^{\log_{10}((30SD_i)^2)}} = \frac{10}{1 + 10^{2(\log_{10}(SD_i) - \log_{10}\left(\frac{1}{30}\right))}}$$
$$= \frac{10}{1 + 10^{2(z - z_0)}}, where\ \mathrm{z} = \log_{10}(SD_i), z_0 = \log_{10}\left(\frac{1}{30}\right)$$

which is a multiplied by a factor of 10 logistic function with the logarithm of *SD* as an argument, centered around the *SD* value of approximately 3% of the cycle. With this formula, (i) extremely stable on- and offsets (with *SD*s smaller than 1% of the cycle) got the highest weights (from approximately 9 to 10), (ii) unstable on- and offsets (with *SD*s larger than 10% of the cycle) got the lowest weights (from 0 to 1), while (ii) rather precise but not extremely precise on- and offsets (with *SD* from 1% to 10% of the cycle) got weights in a range from 1 to 9 (**Supplementary Figure 1A**):

$$w(0.00) = 10, w(0.01) \approx 9, w(0.03) \approx 5, w(0.10) = 1, w(0.30) \approx 0.1$$

Note several properties of the weight function $w(SD)$:

(i) The function $w(SD)$ is monotonically decreasing. It means that smaller weights are assigned to on- and offsets with high variability (less stable modulation) which probably contribute less to precise phasing of the locomotor rhythm.

(ii) The function is limited. The weight tends the supremum of 10 as *SD* tends to 0, and it tends to the infimum of 0 as *SD* tends to infinity.

(iii) The function is non-negative. The infimum is zero thus we effectively filter out the data with extremely high variability.

(iv) The logistic function has a centrally symmetrical shape.

We do not directly use *SD* as the argument for the logistic function but prefer to use $\log_{10}SD$ (the logarithmic scale). The reason for this is the following. The observed *SD* distribution was rather strongly skewed (**Supplementary Figure 1B,D,F**): the population skewness was 0.63 for the onset *SD*, 0.84 for the offset *SD*, and 0.80 for all burst edges (on- and offsets) together. It was possible to make the distribution more symmetrical by using the logarithmic scale (**Supplementary Figure 1C,E,G**): the population skewness was -0.72 for the onset *SD*, -0.53 for the offset *SD*, and -0.61 for all burst edges (on- and offsets) together.

#### 6. Analysis of the population distribution of burst onset and offset phases

We calculated the probability densities $p(\varphi)$ for measured the mean onset $\{\varphi^{on}{}_{i}\}$and offset $\{\varphi^{off}{}_{i}\}$ phases of the $i$'th neuron's burst ($i = 1, \ldots, N$, where $N$ was the number of recorded neurons) with the corresponding standard deviations $\{SD^{on}{}_{i}\}$ and $\{SD^{off}{}_{i}\}$ for the neurons located in spinal segments L4 and L6, respectively. The procedure for a set of $N$ neurons was the following. First, for each $i$'th neuron its phase $\varphi_i$ was represented by a Gaussian distribution with $\sigma = SD_i$ centered at $\varphi_i$:

$$p_i(\varphi) = \frac{1}{\sqrt{2\pi} SD_i} e^{-\frac{(\varphi-\varphi_i)^2}{2SD_i{}^2}}$$

Then, the distributions for phases of individual neurons were averaged across a population:

$$p(\varphi) = \frac{1}{N}\sum_{i=1}^{N} p_i(\varphi) = \frac{1}{N}\sum_{i=1}^{N} \frac{1}{\sqrt{2\pi} SD_i} e^{-\frac{(\varphi-\varphi_i)^2}{2SD_i{}^2}}$$

Such presentation of the phases takes into account variability of the phase in different steps, and also it smoothens the resulting curves.

Analogous to the probability distribution of onset/offset phases, the weighted probability distribution was calculated:

$$p(\varphi) = \frac{\sum_{i=1}^{N} w(SD_i) p_i(\varphi)}{\sum_{i=1}^{N} w(SD_i)} = \frac{\sum_{i=1}^{N} \frac{w(SD_i)}{\sqrt{2\pi} SD_i} e^{-\frac{(\varphi-\varphi_i)^2}{2SD_i{}^2}}}{\sum_{i=1}^{N} w(SD_i)}$$

#### 7. Cluster analysis

In the following cluster analysis, we used squared distance between a phase $\varphi$ and a phase $\psi$: $(\varphi-\psi)^2$. We took into account the circular nature of the phases, so the distance was calculated with a formula:

$$|\varphi-\psi| = \left|\frac{1}{2\pi}Arg\left(e^{2\pi i(\varphi-\psi)}\right)\right|, \qquad (\varphi-\psi)^2 = \left(\frac{1}{2\pi}Arg\left(e^{2\pi i(\varphi-\psi)}\right)\right)^2$$

However, to make our formulae more readable instead of the later long formula, below we wrote briefly $(\varphi-\psi)^2$.

To reveal groups of neurons with similar activity phases, we used a modified version of the "nearest neighbor" method of cluster analysis. Burst phase data were graphed as a scatterplot in which the burst offset phase of a given neuron was plotted *vs* its burst onset (Krouchev et al., 2006) together with the onset and offset weights (**Supplementary Figure 2A**). Each *m*'th cluster consisting of a set of $n_m$ points $\{(\varphi^{on}_i,\varphi^{off}_i)\}$ with the corresponding sets of *SD*s $\{(SD^{on}_i,SD^{off}_i)\}$ and weights $\{(w(SD^{on}_i), w(SD^{off}_i))\}$, $i = 1, \ldots, n_m$, was characterized by its center $(\mathbf{\Phi}^{on}_m,\mathbf{\Phi}^{off}_m)$, estimated as the weighted average of the corresponding phases:

$$\mathbf{\Phi}^{on/off}{}_m = \frac{\sum_{i=1}^{n_m} w(SD^{on/off}{}_i)\, \varphi^{on/off}{}_i}{\sum_{i=1}^{n_m} w(SD^{on/off}{}_i)},$$

and the estimates of *SD*s $(\mathbf{SD}^{on}_m,\mathbf{SD}^{off}_m)$, calculated as the weighted average of the individual *SD*s:

$$\mathbf{SD}^{on/off}{}_m = \frac{\sum_{i=1}^{n_m} w(SD^{on/off}{}_i)\, SD^{on/off}{}_i}{\sum_{i=1}^{n_m} w(SD^{on/off}{}_i)}.$$

The phases $\mathbf{\Phi}^{on}_m$ and $\mathbf{\Phi}^{off}_m$ were called the cluster "burst" on- and offset phases, respectively.
To assess similarity of the *i*'th point $(\varphi^{on}_i,\varphi^{off}_i)$ to the *m*'th cluster $(\mathbf{\Phi}^{on}_m,\mathbf{\Phi}^{off}_m)$, we used two "distance" parameters. One was Euclidian distance $D_{im}$:

$$D_{im} = \sqrt{(\varphi^{on}{}_i - \mathbf{\Phi}^{on}{}_m)^2 + (\varphi^{off}{}_i - \mathbf{\Phi}^{off}{}_m)^2},$$

which was the traditional way to identify bursts with similar mean onset and offset phases. The other parameter was "Z-distance", $Z_{im}$ (the distance calculated after z-score transformation):

$$Z_{im} = \sqrt{\frac{(\varphi^{on}{}_i-\mathbf{\Phi}^{on}{}_m)^2}{SD^{on}{}_i{}^2+\mathbf{SD}^{on}{}_m{}^2} + \frac{(\varphi^{off}{}_i-\mathbf{\Phi}^{off}{}_m)^2}{SD^{off}{}_i{}^2+\mathbf{SD}^{off}{}_m{}^2}},$$

which took into account not only similarity of mean phases but also their variability.

We did not specify the number of clusters *a priori*. The clustering procedure consisted of three stages: (i) ranking neurons according to stability of their activity, (ii) primary clustering ("seeding") that produced an initial set of "raw" clusters, (iii) "refining" of the "raw" clusters.

First, the set of points $\{(\varphi^{on}_i,\varphi^{off}_i)\}$ with the corresponding *SD*s $\{(SD^{on}_i,SD^{off}_i)\}$ was sorted in the descending order according to their integral stability [sum of squared weights $w^2(SD^{on}_i)$ + $w^2(SD^{off}_i)$]. This ranked list was used for further analysis.

Then, for "seeding", each point $(\varphi^{on}_i,\varphi^{off}_i)$ one by one was compared with already existing clusters with 2 different outcomes: (i) if the point $(\varphi^{on}_i,\varphi^{off}_i)$ with *SD*s $(SD^{on}_i,SD^{off}_i)$ was far from all already existing *m* clusters (both $D_{ik} > D_{max}$ and $Z_{ik} > Z_{max}$, for all $k = 1, \ldots, m$; for "seeding" we used $D_{max} = 0.15$, $Z_{max} = 3$) then it formed a new $(m+1)$'th cluster with $(\mathbf{\Phi}^{on}_m,\mathbf{\Phi}^{off}_m) = (\varphi^{on}_i,\varphi^{off}_i)$ and $(\mathbf{SD}^{on}_m,\mathbf{SD}^{off}_m) = (SD^{on}_i,SD^{off}_i)$ (the first point in the list always formed the first cluster); (ii) if the point was close to one or several different clusters then it was added to the cluster for which $Z_{im}$ was minimal, and the cluster parameters $(\mathbf{\Phi}^{on}_m,\mathbf{\Phi}^{off}_m)$ and $(\mathbf{SD}^{on}_m,\mathbf{SD}^{off}_m)$ were updated, (iii) then the next point of the set was considered and so on.

The set of clusters obtained during the "seeding" stage was used to resort the points with a "refining" procedure, similar to "seeding". Specifically, each point $(\varphi^{on}_i,\varphi^{off}_i)$ was compared with already existing clusters with 2 different outcomes: (i) if the point $(\varphi^{on}_i,\varphi^{off}_i)$ with *SD*s $(SD^{on}_i,SD^{off}_i)$ was far from all already existing *m* clusters ($D_{ik} > D_{max}$ and $Z_{ik} > Z_{max}$, for all $k = 1, \ldots, m$; for "refining" we used $D_{max} = 0.1$, $Z_{max} = 3$) then it was added to "assorted" (*Cluster* #0); (ii) if the point was close to one or several different clusters, then it was added to the cluster for which $D_{im}$ was minimal, and the cluster parameters $(\mathbf{\Phi}^{on}_m,\mathbf{\Phi}^{off}_m)$ and $(\mathbf{SD}^{on}_m,\mathbf{SD}^{off}_m)$ were updated. After the resorting, clusters containing too few neurons ($\leq 2$) were deleted, and their neurons were added to “assorted” (*Cluster* #0). This resorting was repeated until convergence to the final set of clusters (**Supplementary Figure 2**). The heatmap matrixes for the Euclidean distance and Z-distance illustrating the results of clustering are presented in **Supplementary Figure 3, A** and **B**, respectively. The main reason for using Z-distance in addition to Euclidian distance was to avoid fusing well-defined neighboring clusters with very small ($\mathbf{SD}^{on}$,$\mathbf{SD}^{off}$) (e.g., #1 and #2 or #6 and #8 in **Figure 5**) despite the small differences in the mean phases ($\mathbf{\Phi}^{on}$,$\mathbf{\Phi}^{off}$). We tried a simplified version of the clustering procedure without Z-distance and obtained qualitatively similar but quantitatively worse results: (i) more neurons remained assorted (19% without Z-distance compared to only 9% with Z-distance), (ii) some clusters (#12 and #13) were not revealed, and (iii) some neighboring clusters became fused (#5 with #11, #2 with #12, #6 with #8 and #16).

There were several reasons for our choice of the threshold value $D_{max}$. First, the swing comprised 0.38 of the entire cycle. Therefore, the $D_{max}$ = 0.1 that we used guaranteed that all bursts of all neurons in a *Cluster* were similar. The looser criteria (e.g., $D_{max}$ = 0.2) might have mixed in the same *Cluster* neurons with functionally different burst phases. On the other hand, the stricter criteria (e.g., $D_{max}$ = 0.09, 0.08, 0.07, 0.06) yielded *Clusters* with practically the same mean values ($\mathbf{\Phi}^{on}$,$\mathbf{\Phi}^{off}$), but many points remained assorted (14%, 17%, 24%, 38%, correspondingly; data not shown). The same reasons determined our choice of the threshold value $Z_{max}$.

#### 8. Combining Clusters into Subgroups

Among 19 *Clusters* that were revealed by the cluster analysis, there were pairs of *Clusters* which midburst phases were almost exactly in anti-phase (*Clusters* #2 and #5, #8 and #13, #18 and #19, etc.; **Supplementary Figure 4A**). Existence of such anti-phase pairs was not guaranteed *a priori*: in a hypothetical example presented in **Supplementary Figure 4B** there are no anti-phase pairs. On the other hand, as the locomotor movements consist of reciprocal rhythmic flexions/extensions, the existence of subpopulations of neurons active in anti-phase are not surprising.

To look for such cases, for each pair of *Clusters* $i$ and $j$ with on/offest phases ($\mathbf{\Phi}^{on}{}_i$, $\mathbf{\Phi}^{off}{}_i$) and ($\mathbf{\Phi}^{on}{}_j$, $\mathbf{\Phi}^{off}{}_j$) we calculated the midburst phases $\mathbf{\Phi}^{mid}{}_i$ and $\mathbf{\Phi}^{mid}{}_j$, then the distance between the midburst phases, and finally the deviation of this difference from 0.5 (deviation from the exact anti-phase activity $a_{ij}$ ("the anti-phase index for a pair of *Subgroups*"):

$$a_{ij} = 0.5 - \left|\mathbf{\Phi}^{mid}{}_i - \mathbf{\Phi}^{mid}{}_j\right|$$

An example of such calculations for *Cluster* #13 *vs* all other *Clusters* is shown in **Supplementary Figure 4C**. *Clusters* #13 and #8 are almost exactly in anti-phase: $a_{13,8} < 0.01$. *Clusters* #13 and #6,9 are close to anti-phase: $a_{13,6} < 0.03$, $a_{13,9} < 0.03$. *Clusters* #13 and #16 are not far from anti-phase: $a_{13,16} < 0.06$. Burst of all other *Clusters* are far from the anti-phase relationship with *Cluster* #13: $a_{13,i} > 0.10$.

Results of these calculations for all observed pairs of *Clusters* are presented in **Supplementary Figure 5D-F** (the same results are shown in **Figure 6A**). One can notice that all 19 *Clusters* can be combined in 4 pairs of *Subgroups* (totally 8 *Subgroups*; borders between *Subgroups* are shown with white dashed lines in **Supplementary Figure 5A,D**) so that *Clusters* within one pair the *Subgroups* are approximately in anti-phase: *Clusters* ##10,13,17 from *Subgroup* #1 are approximately in anti-phase with *Clusters* ##6,8,9,16 from *Subgroup* #2, *Clusters* from *Subgroup* #3 are approximately in anti-phase with *Clusters* from *Subgroup* #4, and so on.

To evaluate whether this grouping is the optimal one, we tested all other possible combinations of *Clusters* into reciprocal pairs of *Subgroups*. Let us consider $2N$ *Subgroups* $\{S_1, S_2, \ldots, S_{2N-1}, S_{2N}\}$ making up a combination **C** consisting of $N$ pairs $\mathbf{C}=\{(S_1, S_2), \ldots, (S_{2N-1}, S_{2N})\}$ so that in the $k$'th pair *Subgroup* $S_{2k-1}$ is approximately in anti-phase with *Subgroup* $S_{2k}$. (In an example shown in **Supplementary Figure 5B,E** there are 3 pairs of *Subgroups*: *Subgroup* #1 is approximately in anti-phase with *Subgroup* #2, *Subgroup* #3 is approximately in anti-phase with *Subgroup* #4, *Subgroup* #5 – with *Subgroup* #6.) Each *Subgroup* consists of *Clusters*, therefore for the $k$'th pair of *Subgroups* there is the corresponding set of *Cluster* pairs $P_k$:

$$P_k = \{(i,j) | i \in S_{2k-1}, j \in S_{2k}\}$$

with the size of this set $|P_k|$. (In an example shown in **Supplementary Figure 5B,E** for the 2nd pair of *Subgroups*, consisting of *Subgroups* ##3,4, the set of *Cluster* pairs $P_2=\{(18, 4), (18, 19)\}$.)

Then for the selected set of *Subgroup* pairs one can calculate the *selected* cost function:

$$selected(\mathbf{C}) = \frac{\sum_{k=1}^{N} \sum_{(i,j)\in P_k} a_{ij}}{\sum_{k=1}^{N} |P_k|}$$

which is, obviously, equal to the average value of the parameter $a_{ij}$ for all pairs of *Clusters* in the combination **C**. The closer the *selected* (**C**) to 0, the better we combined *Clusters* into reciprocally active pairs. By other words, in an example shown in **Supplementary Figure 5E** one gets the average value of the parameter $a_{ij}$ for all cells demarcated by the red, green, and dark blue dashed rectangles.

Then one can consider all possible combinations **C** consisting of $2N$ *Subgroups* and find the combination $\mathbf{C_{min}}$ for which the *selected* cost is minimal. This combination $\mathbf{C_{min}}$ we call "the optimal combination of $N$ pairs of *Subgroups*".

(Note that one can consider also pairs ($S_2$, $S_1$), …, ($S_{2N}$, $S_{2N-1}$). However, as the matrix is symmetric: $a_{ij} = a_{ji}$ – this would not change the result.)

With an increase in *N*, this *selected* cost is mostly decreasing (red line in **Supplementary Figure 6A**). The minimum is reached at 8 pairs of *Subgroups*, consisting totally of 16 *Subgroups* (this optimal combination of 8 pairs is presented in **Supplementary Figure 5C,F**). For this combination almost all *Subgroups* include just one *Cluster*. *Clusters* in the opposing *Subgroups* are almost precisely in anti-phase (e.g. *Clusters* ##13,17 belonging to *Subgroup* #3 are almost precisely in anti-phase with *Clusters* ##8,9 belonging to *Subgroup* #4).

However, one can easily see a weakness in this combination of *Clusters*: often a pair of *Clusters* that is almost in anti-phase, is not included in any of the $P_k$ (e.g. pairs (13,16), (13,6), (17,16), (17,6) in **Supplementary Figure 5F**, are outside of the demarcated areas). This result looks like over-fitting. To evaluate the degree of such overfitting, for the optimal combination $\mathbf{C_{optimal}}$ one can consider the set $N_k$ that includes all "neighbor" to the selected pairs of *Clusters* (marked by black crosses in **Supplementary Figure 5E,F**):

$$N_k = \{(i,j) | i \in S_{2k-1}, j \in S_{2k+2}\} \cup \{(i,j) | i \in S_{2k+1}, j \in S_{2k}\}$$

(Obviously, to take into account the circular nature of the data, one should set $S_{2N+1}=S_1$, $S_{2N+2}=S_2$.) And then the corresponding *neighbor* cost function can be defined as:

$$neighbor(\mathbf{C_{optimal}}) = \frac{\sum_{k=1}^{N} \sum_{(i,j)\in N_k} a_{ij}}{\sum_{k=1}^{N} |N_k|}$$

With an increase in *N* this *neighbor* cost is decreasing (blue line in **Supplementary Figure 6A**). It means that overfitting gets worse. To find an optimal combination of *Subgroups*, we considered "the contrast ratio", *contrast*:

$$contrast = \frac{neighbor(\mathbf{C_{optimal}})}{selected(\mathbf{C_{optimal}})}$$

As one can see in **Supplementary Figure 6B**, the maximum of the *contrast* is reached at 4 pairs of *Subgroups*. These 4 pairs are exactly the *Subgroups* presented in **Supplementary Figure 5A,D**. With this combination the *selected* cost is 0.03: the corresponding *Clusters* are active in anti-phase with an average precision of just 3% of the locomotor cycle. On the other hand, the *neighbor* cost is 0.13: the corresponding neighbor *Clusters* are not in anti-phase with an average precision of 13% of the locomotor cycle. Therefore, we think that our combination of *Clusters* into 4 pairs of *Subgroups* looks reasonably good.

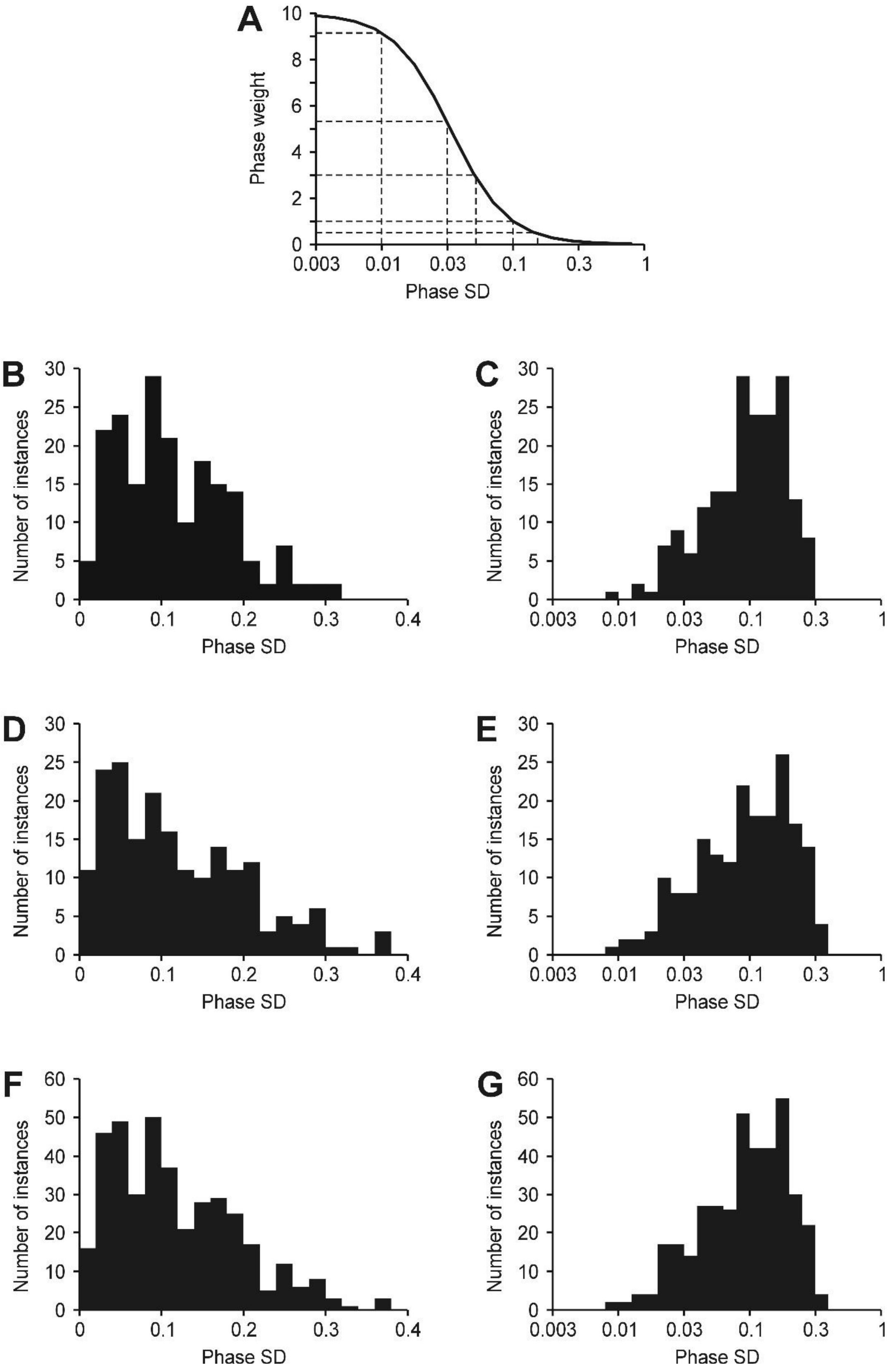


**Supplementary Figure 1.** (**A**) The phase weight *vs* the phase SD curve. (**B-G**) Distributions of phase SDs for onsets (**B,C**), offsets (**D,E**), and edges (onsets and offsets together) (**F,G**), respectively, in the entire population of active modulated neurons. Horizontal scale in **A,C,E,G** is logarithmic.

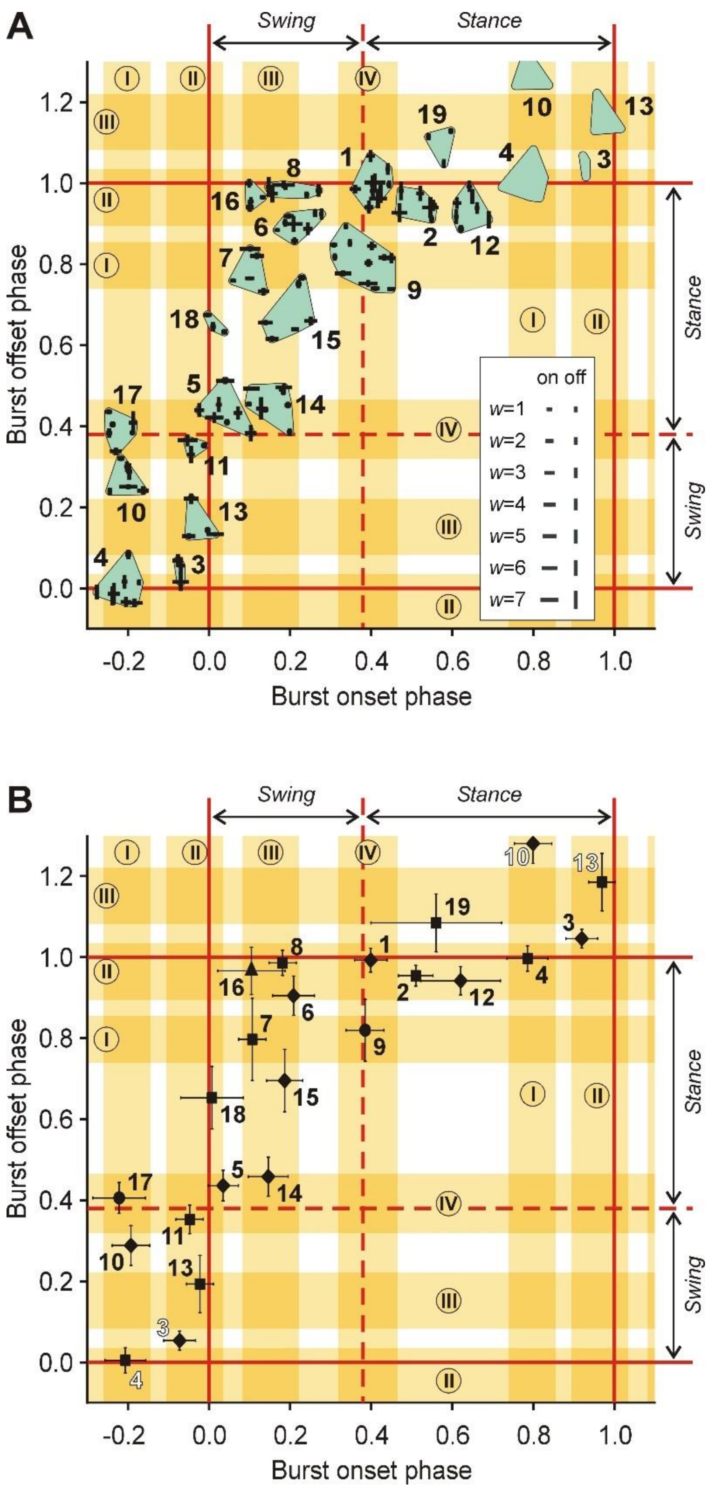


**Supplementary Figure 2.** Activity phases of 126 stable neurons and their *Clusters*. (**A**) Results of clustering of the stable neurons into 19 *Clusters*. Lengths of the horizontal and vertical arms of the crosses indicate weights of the burst onset and offset of individual neurons, respectively. Green areas demarcate *Clusters*. (**B**) Scatterplot for average (±*SD*) *Cluster* offset phase plotted *vs* onset phase. These two panels present the same data as **Figure 5** but in a two-dimensional form that was used for cluster analysis of motor activity by other authors (Krouchev et al., 2006). Four phase ranges I-IV (indicated by yellow) are the same as in **Figure 5**. Note that due to the clustering procedure, *Clusters* with the smaller numbers tend to have smaller SDs, while *Clusters* with the greater numbers tend to have larger SDs.

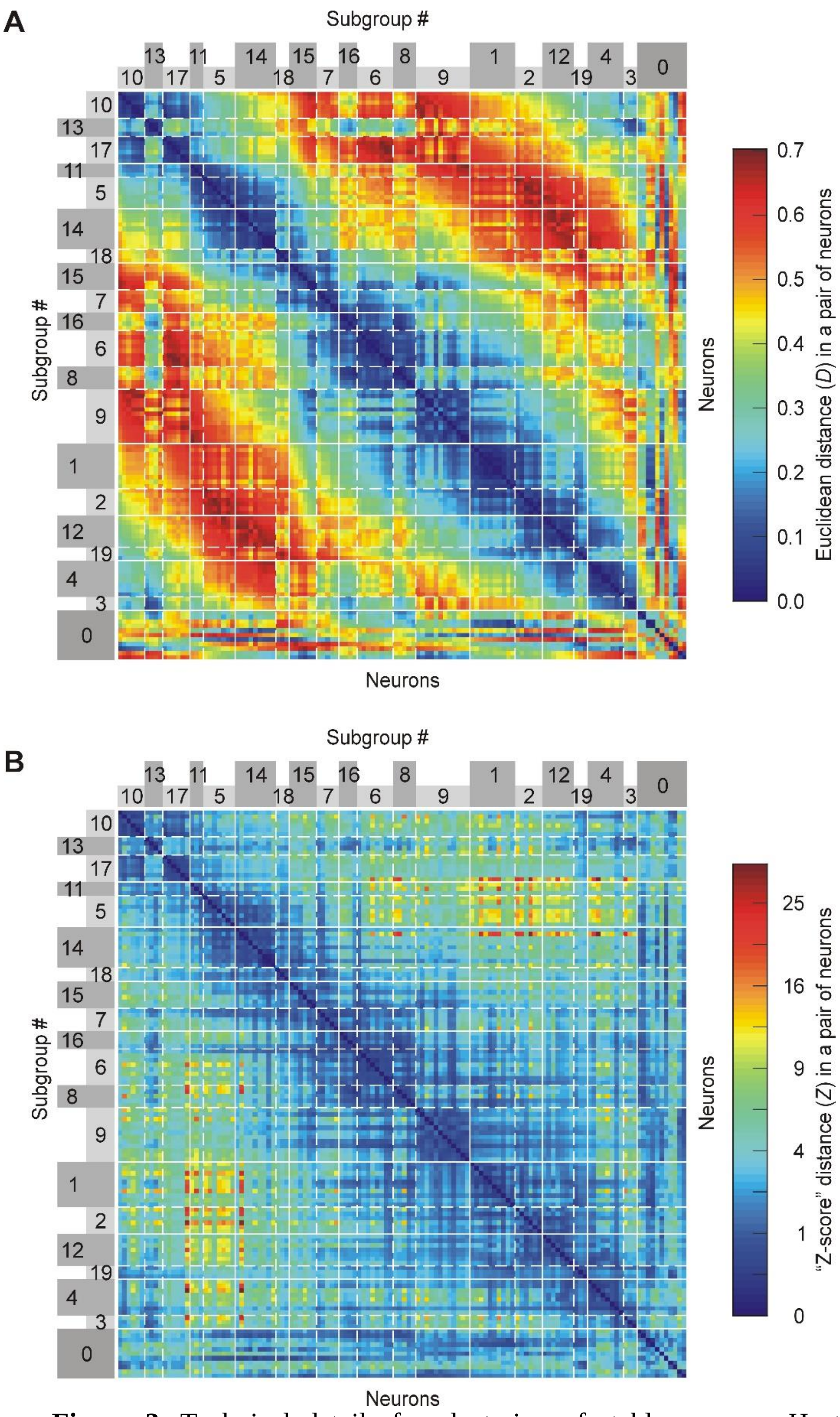


**Supplementary Figure 3.** Technical details for clustering of stable neurons. Heatmaps of the Euclidean (**A**) and "z-score" (**B**) distances for all pairs of 126 clustered neurons. Our analysis produced 19 *Clusters*, while 9% of the neurons were not included into any of the 19 *Clusters* and were called "*Cluster* #0". Note that for any pair of neurons belonging to the same *Cluster*, Euclidean distance was <0.1 (dark blue squares on the diagonal in **A**) and "z-score" distances was <3 (dark blue squares on the diagonal in **B**). Note the non-linear scale in (**B**), it was used to have better color resolution for small values of Z. Numbers were assigned to *Clusters* according to the clustering procedure.

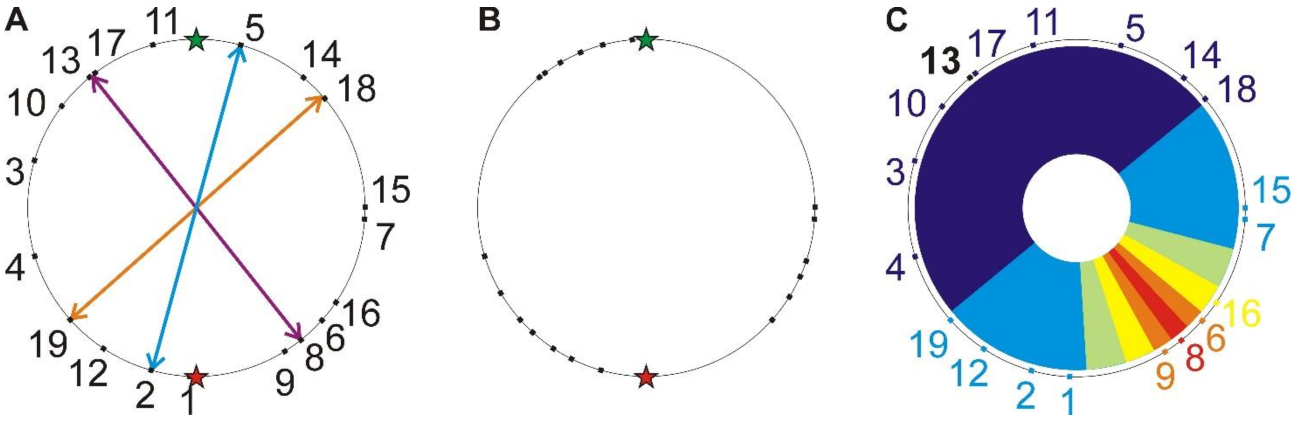


**Supplementary Figure 4.** Search for *Clusters* active approximately in anti-phase. (**A**) A circular diagram showing phases of the middle of the burst for all 19 revealed *Clusters* on the locomotor cycle circle. Some *Clusters* were active in anti-phase with one another, e.g.: *Clusters* #2 and #5 (blue arrow), *Clusters* #8 and #13 (purple arrow), *Clusters* #18 and #19 (orange arrow). (**B**) A circular diagram showing the mid-burst phases of the *a priori* possible hypothetical *Clusters* neither of which are active in anti-phase. In **A** and **B** the green and red stars indicated the middle of the swing and stance, respectively. (**C**) An example of phase differences between a reference *Cluster* (#13, shown by bold black) and the other *Clusters*. The color scheme is the same as in **Figure 6A**: the difference from anti-phase less than 1% of the cycle duration is indicated by red, <3% – by orange, <6% – by yellow, <10% – by green, <25% – by light blue, the in-phase half of the cycle is shown by dark blue.

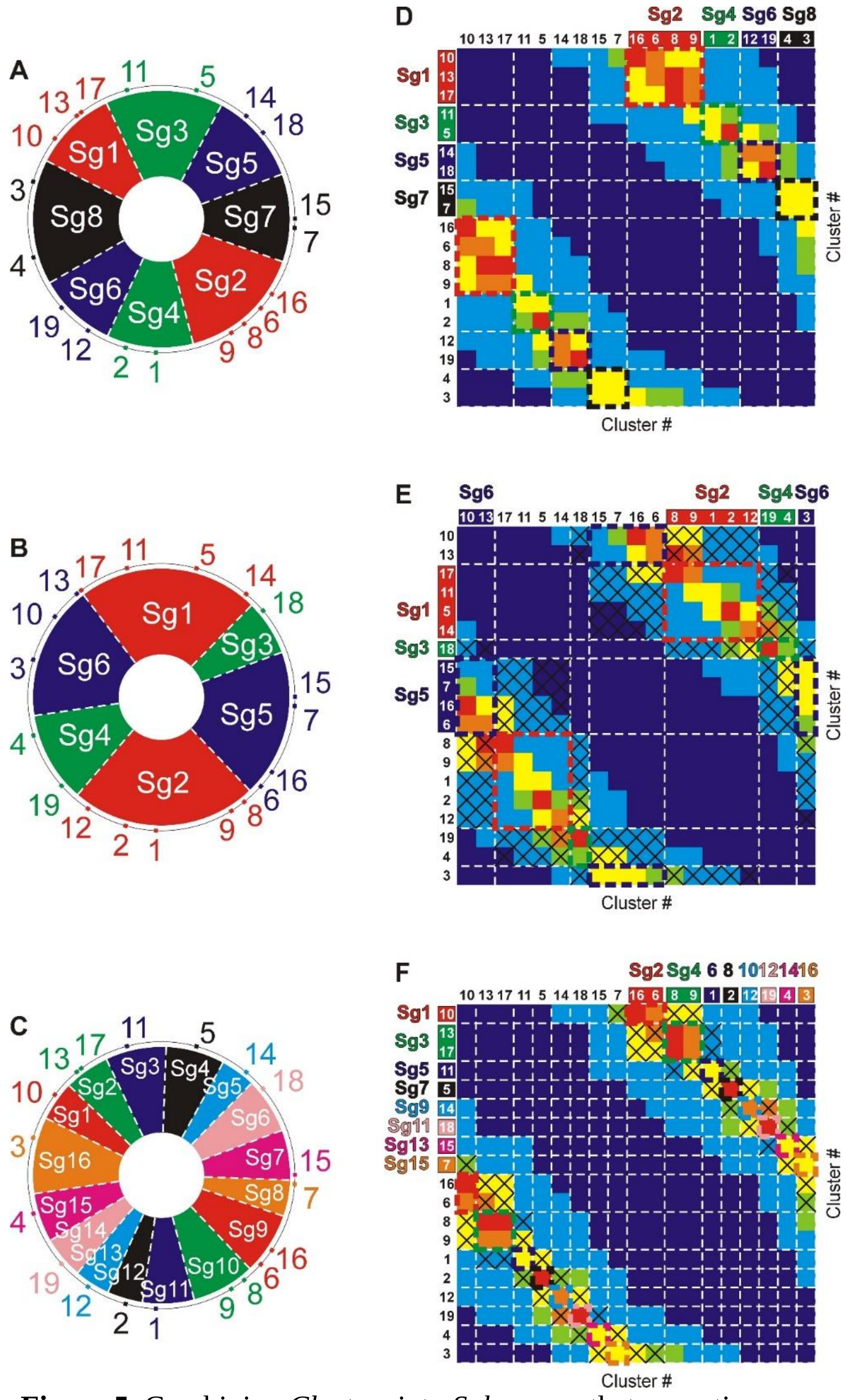


**Supplementary Figure 5.** Combining *Clusters* into *Subgroups* that are active approximately in anti-phase. (**A-C**) All *Clusters* were combined into *N* pairs of *Subgroups*: a combination with 4 pairs (**A**), a combination with 3 pairs (**B**), a combination with 8 pairs (**C**). The midphase of each *Cluster* is shown on the locomotor cycle circle. Borders between *Subgroups* are shown with white dashed lines. *Subgroups* active in anti-phase are shown with the same color (e.g., in **A** *Subgroups* ##1,2 are shown in red, *Subgroups* ##3,4 – in green, *Subgroups* ##5,6 – in blue, *Subgroups* ##7,8 – in black). (**D-F**) The heatmap of the phase shift between midbursts of different *Cluster* pairs corresponding to different *Cluster* combinations: **D** corresponds to 4 pairs of *Subgroups* shown in **A**, **E** – 3 pairs of *Subgroups* shown in **B**, and **F** – 8 pairs of *Subgroups* shown in **C**. White dashed lines show borders between *Subgroups*. Dashed color rectangles demarcate “selected” *Cluster* pairs. Black crosses in (**E,F**) indicate “neighbor” *Cluster* pairs. The color scheme is the same as in **Figure 6A** and **Supplementary Figure 4C**.

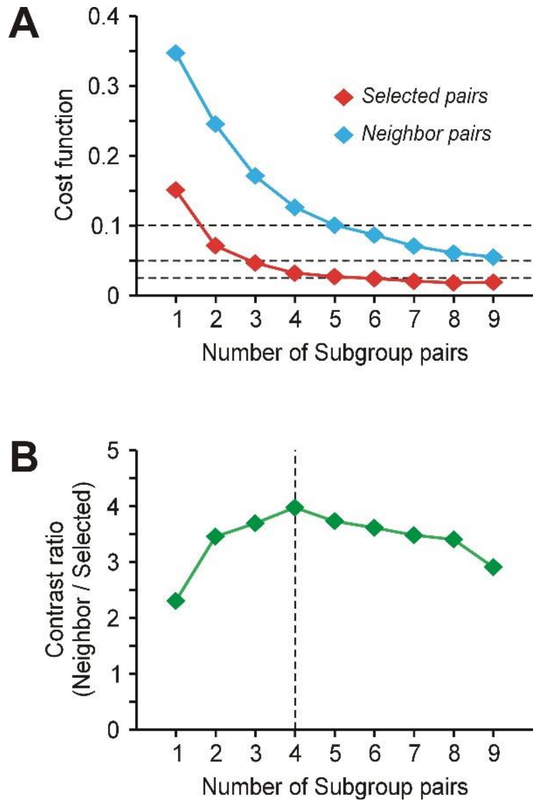


**Supplementary Figure 6.** (**A**) The *selected* cost for the optimal combinations of *Clusters* (the minimum of the *selected* cost function across all possible combinations of *Clusters* for a given number of *Subgroup* pairs *N*) is shown in red. The corresponding *neighbor* cost for these optimal combinations is shown in blue. (**B**) The contrast ratio (the *neighbor* cost divided by the *selected* cost for the optimal combination of *Clusters* into *N* pairs of *Subgroups*). The maximum is reached at *N*=4. The optimal combination $\mathbf{C_{optimal}}$ of 4 *Cluster* pairs is presented in **Supplementary Figure 5A,D**.

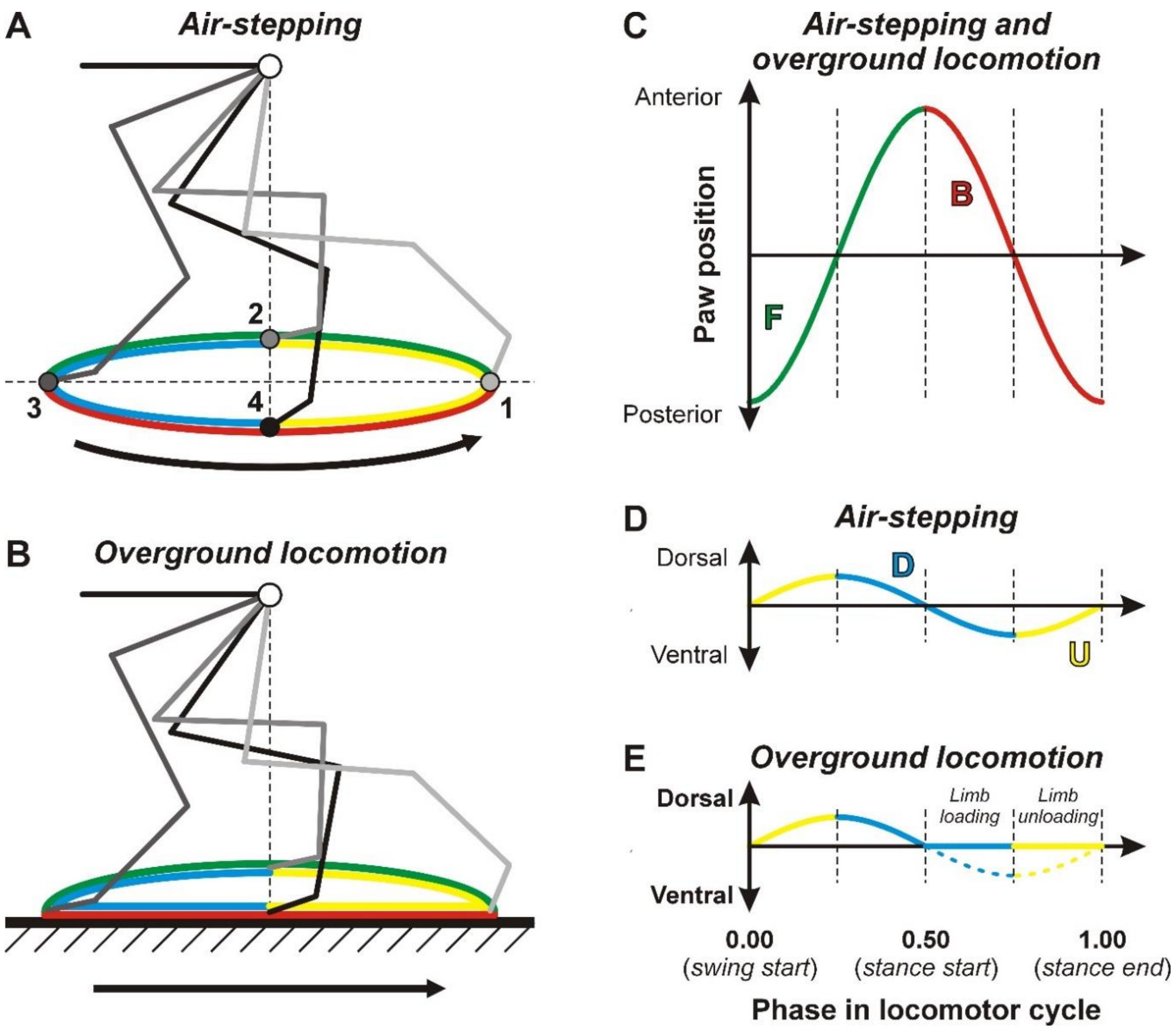


**Supplementary Figure 7.** Vertical and horizontal components of the locomotor movements. During air-stepping, the hindlimb movements can be presented with an idealized scheme (**A**). The paw trajectory has 4 critical points: 1) the extreme posterior (light gray limb configuration), 2) the extreme dorsal (gray limb configuration), 3) the extreme anterior (dark gray limb configuration), 4) the extreme ventral (black limb configuration). These critical points divide the entire limb trajectory into forward (green, F; from point 1 to point 3) and backward (red, B; from point 3 to point 1) movements, or into upward (yellow, U; from point 4 to point 2) and downward (blue, D; from point 2 to point 4) movements. A black arrow indicates the direction of movement. The corresponding graphs for phase dependence of the anteroposterior (**C**) and dorso-ventral (**D**) paw position are drawn under assumption of equal swing and stance durations. Analogous to (**A**), a scheme of the hindlimb movements can be drawn for overground locomotion (**B**). In contrast to (**A**), the lower part of the paw trajectory in (**B**) is horizontal (the limb is on the ground during stance). However, it is not uniform in terms of ground reaction force vertical component: in the red-blue part of the trajectory, gradual loading of the limb takes place, while in the red-yellow part, the limb gets unloaded. The phase dependence of the antero-posterior paw position for overground locomotion is the same as for the air stepping (**C**). The phase dependence of the dorso-ventral paw position is slightly different: during stance the position does not change, while the force does (**E**; solid and dashed lines, respectively). Despite the difference in kinematics in forces, one can argue that the red-blue part of the trajectory during air stepping (when the paw moves downwards due to activity of the extensor muscles) is homologous to the red-blue part of the trajectory during overground locomotion (when the same extensor muscles are activated to counteract the increasing load). Similarly, the red-yellow part of the air stepping trajectory (when the paw moves upwards due to activation of the flexors and activity reduction of the extensors) is homologous to the red-yellow part of the overground trajectory (when the limb unloading coincides with similar changes in the muscle activity).
